# Parallel loss of symbiosis genes in relatives of nitrogen-fixing non-legume *Parasponia*

**DOI:** 10.1101/169706

**Authors:** Robin van Velzen, Rens Holmer, Fengjiao Bu, Luuk Rutten, Arjan van Zeijl, Wei Liu, Luca Santuari, Qingqin Cao, Trupti Sharma, Defeng Shen, Yuda P. Roswanjaya, Titis A.K. Wardhani, Maryam Seifi Kalhor, Joëlle Jansen, D. Johan van den Hoogen, Berivan Güngör, Marijke Hartog, Jan Hontelez, Jan Verver, Wei-Cai Yang, Elio Schijlen, Rimi Repin, Menno Schilthuizen, M. Eric Schranz, Renze Heidstra, Kana Miyata, Elena Fedorova, Wouter Kohlen, Ton Bisseling, Sandra Smit, Rene Geurts

## Abstract

Rhizobium nitrogen-fixing nodules are a well-known trait of legumes, but nodules also occur in other plant lineages either with rhizobium or the actinomycete *Frankia* as microsymbiont. The widely accepted hypothesis is that nodulation evolved independently multiple times, with only a few losses. However, insight in the evolutionary trajectory of nodulation is lacking. We conducted comparative studies using *Parasponia* (Cannabaceae), the only non-legume able to establish nitrogen fixing nodules with rhizobium. This revealed that *Parasponia* and legumes utilize a large set of orthologous symbiosis genes. Comparing genomes of *Parasponia* and its non-nodulating relative *Trema* did not reveal specific gene duplications that could explain a recent gain of nodulation in *Parasponia*. Rather, *Trema* and other non-nodulating species in the Order Rosales show evidence of pseudogenization or loss of key symbiosis genes. This demonstrates that these species have lost the potential to nodulate. This finding challenges a long-standing hypothesis on evolution of nitrogen-fixing symbioses, and has profound implications for translational approaches aimed at engineering nitrogen-fixing nodules in crop plants.

## Introduction

Nitrogen sources such as nitrate or ammonia are key nutrients for plant growth, but their availability is frequently limited. Some plant species in the related orders Fabales, Fagales, Rosales and Cucurbitales-collectively known as the nitrogen fixation clade-can overcome this limitation by establishing a nitrogen-fixing endosymbiosis with either *Frankia* or rhizobium bacteria^1^. These symbioses require specialized root organs, known as nodules, that provide optimal physiological conditions for nitrogen fixation^2^. For example, nodules of legumes (Fabaceae, order Fabales) contain a high concentration of hemoglobin that is essential to control oxygen homeostasis and protect the rhizobial nitrogenase enzyme complex from oxidation^2,3^. Legumes, such as soybean (*Glycine max*) and common bean (*Phaseolus vulgaris*), represent the only crops that possess nitrogen-fixing nodules, and engineering this trait in other crop plants is a long-term vision in sustainable agriculture^4,5^.

Nodulating plants represent ~10 clades that diverged >100 million years ago and which are nested in many non-nodulating lineages^6,7^. Consequently, the widely accepted hypothesis is that nodulation originated independently multiple times, preceded by a shared hypothetical predisposition event in a common ancestor of the nitrogen fixation clade^1,6–9^. Genetic dissection of rhizobium symbiosis in two legume models-*Medicago truncatula* (medicago) and *Lotus japonicus* (lotus)-has uncovered symbiosis genes that are essential for nodule organogenesis, bacterial infection, and nitrogen fixation (Supplementary Table 1). These include genes encoding LysM-type receptors that perceive rhizobial lipo-chitooligosaccharides (LCOs) and transcriptionally activate the *NODULE INCEPTION* (*NIN*) transcription factor^10–15^. Expression of *NIN* is essential and sufficient to set in motion nodule organogenesis^14,16–18^. Some symbiosis genes have been co-opted from the more ancient and widespread arbuscular mycorrhizae symbiosis^19,20^. However, causal genetic differences between nodulating and non-nodulating species have not been identified^21^.

To obtain insight in the evolution of rhizobium symbiosis we conducted comparative studies using *Parasponia* (Cannabaceae, order Rosales). The genus *Parasponia* is the only lineage outside the legume family establishing a nodule symbiosis with rhizobium^22–25^. Similar as shown for legumes, nodule formation in *Parasponia* is initiated by rhizobium-secreted LCOs and this involves a close homolog of the legume LysM-type receptor NOD FACTOR PERCEPTION / NOD FACTOR RECEPTOR 5 (NFP/NFR5)^26–28^. This suggests that *Parasponia* and legumes utilize a similar set of genes to control nodulation. The genus *Parasponia* represents a clade of five species that is phylogenetically embedded in the closely related *Trema* genus^29^. Like *Parasponia* and most other land plants, *Trema* species can establish an arbuscular mycorrhizae symbiosis (Supplementary Fig. 1)., However, they are non-responsive to rhizobium LCOs and do not form nodules^25,28^. Taken together, *Parasponia* is an excellent system for comparative studies with legumes and non-nodulating *Trema* species to provide insights into the evolutionary trajectory of nitrogen-fixing root nodules.

## RESULTS

### Nodule organogenesis is a dominant genetic trait

First, we took a genetics approach for understanding the rhizobium symbiosis trait of *Parasponia* by making intergeneric crosses (Supplementary Table 2). Viable F_1_ hybrid plants were obtained only from the cross *Parasponia andersonii* (2n=20) × *Trema tomentosa* (2n=4×=40) (Fig. 1a, Supplementary Fig. 2). These triploid hybrids (2n=3×=30) were infertile, but could be propagated clonally. We noted that F_1_ hybrid plants formed root nodules when grown in potting soil, similar as earlier observations for *P. andersonii^30^*. To further investigate the nodulation phenotype of these hybrid plants, clonally propagated plants were inoculated with two different strains; *Bradyrhizobium elkanii* strain WUR3^30^ or *Mesorhizobium plurifarium* strain BOR2. The latter strain was isolated from the rhizosphere of *Trema orientalis* in Malaysian Borneo and showed to be an effective nodulator of *P. andersonii* (Supplementary Fig. 3). Both strains induced nodules on F_1_ hybrid plants (Fig. 1b,d,e; Supplementary Fig. 4) but, as expected, not on *T. tomentosa*, nor on any other *Trema* species investigated. Using an acetylene reduction assay we noted that, in contrast to *P. andersonii* nodules, in F_1_ hybrid nodules of plant H9 infected with *M. plurifarium* BOR2 there is no nitrogenase activity (Fig. 1c). To further examine this discrepancy, we studied the cytoarchitecture of these nodules. In *P. andersonii* nodules, apoplastic *M. plurifarium* BOR2 colonies infect cells to form so-called fixation threads (Fig. 1f,h-j), whereas in F_1_ hybrid nodules these colonies remain apoplastic, and fail to establish intracellular infections (Fig. 1g,k). To exclude the possibility that the lack of intracellular infection is caused by heterozygosity of *P. andersonii* where only a nonfunctional allele was transmitted to the F_1_ hybrid genotype, or by the particular rhizobium strain used for this experiment, we examined five independent F_1_ hybrid plants either inoculated with *M. plurifarium* BOR2 or *B. elkanii* WUR3. This revealed a lack of intracellular infection structures in nodules of all F_1_ hybrid plants tested, irrespective which of both rhizobium strains was used (Fig. 1g,k, Supplementary Fig. 4), confirming that heterozygosity of *P. andersonii* does not play a role in the F_1_ hybrid infection phenotype. These results suggest, at least partly, independent genetic control of nodule organogenesis and rhizobium infection. Since F_1_ hybrids are nodulated with similar efficiency as *P. andersonii* (Fig. 1b), we conclude that the network controlling nodule organogenesis is genetically dominant.

**Figure 1:**
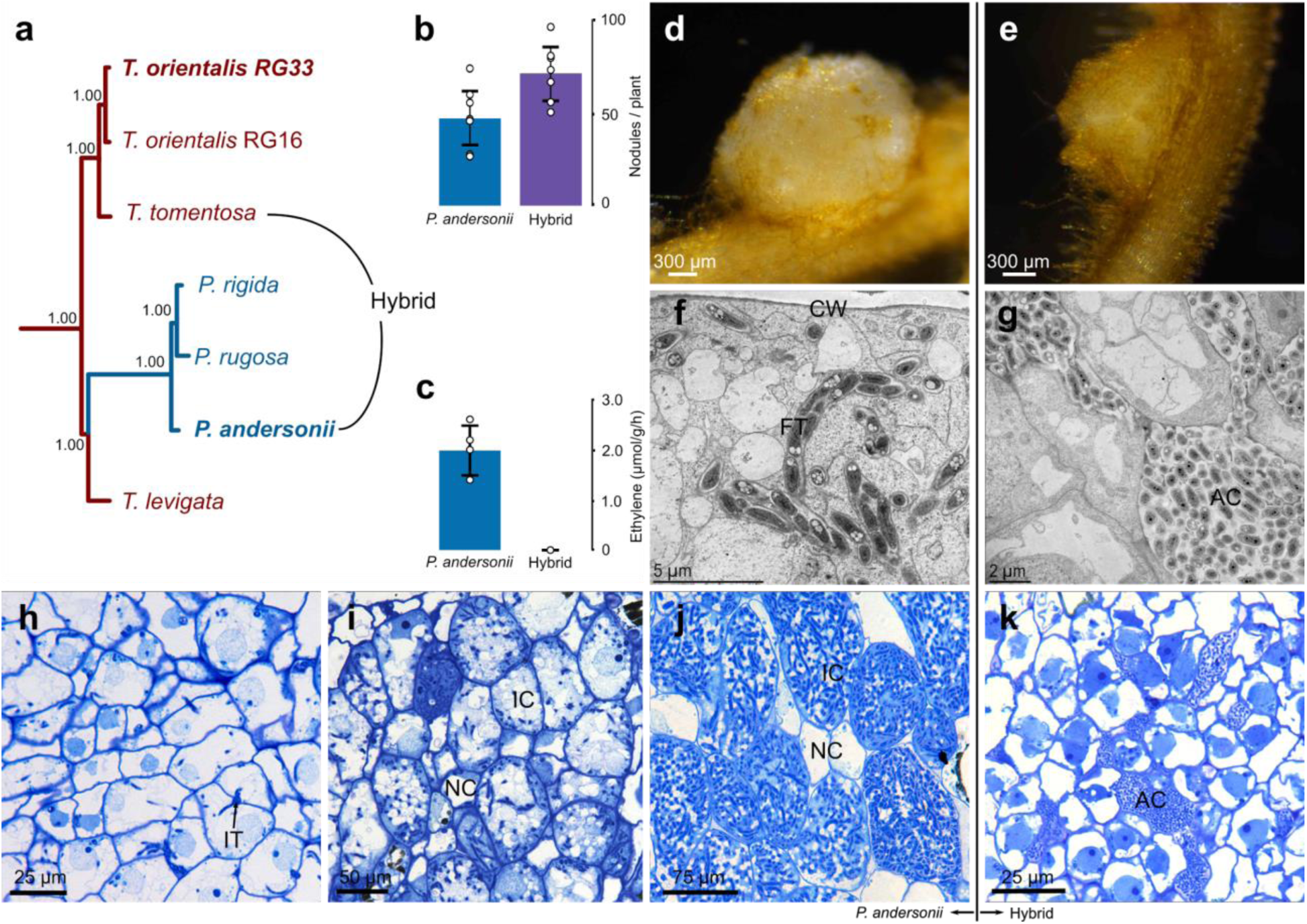
Nodulation phenotype of *Parasponia* and interspecific *Parasponia* × *Trema* F_1_ hybrid plants. **(a)** Phylogenetic reconstruction based on whole chloroplast of *Parasponia* and *Trema*. The *Parasponia* lineage (marked blue) is embedded in the *Trema* genus (marked red). Species selected for interspecific crosses are indicated, species used for reference genome assembly are in bold. Node labels indicate posterior probabilities. **(b)** Mean number of nodules on roots of *P. andersonii* and *P. andersonii* × *T. tomentosa* F_1_ hybrid plants (n=7). **(c)** Mean nitrogenase activity in acetylene reductase assay of *P. andersonii* and *P. andersonii* × *T. tomentosa* F_1_ hybrid nodules (n=4). Barplot error bars indicate standard deviations; dots represent individual measurements **(d)** *P. andersonii* nodule. **(e)** *P. andersonii* × *T. tomentosa* F_1_ hybrid nodule. **(f,g)** Ultrastructure of nodule tissue of *P. andersonii* **(f)** and F_1_ hybrid **(g)**. Note the intracellular fixation thread (FT) in the cell of *P. andersonii* in comparison with the extracellular, apoplastic colonies of rhizobia (AC) in the hybrid nodule. **(h-i)** Light microscopy images of *P. andersonii* nodules in three subsequent developmental stages. **(h)** Stage 1: initial stages of colonization when infection threads (IT) enter the host cells. **(i)** Stage 2: progression of rhizobium infection in nodule host cell, **(j)** Stage 3: nodule cells completely filled with fixation threads. Note difference in size between the infected (IC) and non-infected cells (NC). **(k)** Light microscopy image of F1 hybrid nodule cells. Note rhizobium colonies in apoplast, surrounding the host cells (AC). Nodules have been analysed 6 weeks post inoculation with *Mesorhizobium plurifarium* BOR2. Abbreviations: FT: fixation thread, CW: cell wall, AC: apoplastic colony of rhizobia, IT: infection threads, IC: infected cell, NC: non-infected cell.

### *Parasponia* and *Trema* genomes are highly similar

Based on preliminary genome size estimates using FACS measurements, three *Parasponia* and five *Trema* species were selected for comparative genome analysis (Supplementary Table 3). K-mer analysis of medium-coverage genome sequence data (~30x) revealed that all genomes had low levels of heterozygosity, except those of *Trema levigata* and *T. orientalis* (accession RG16) (Supplementary Fig. 5). Based on these k-mer data we also generated more accurate estimates of genome sizes. Additionally, we used these data to assemble chloroplast genomes based on which we obtained additional phylogenetic evidence that *T. levigata* is sister to *Parasponia* (Fig. 1a, Supplementary Fig. 6–8). Graph-based clustering of repetitive elements in the genomes (calibrated with the genome size estimates based on k-mers) revealed that all selected species contain roughly 300 Mb of non-repetitive sequence, and a variable repeat content that correlates with the estimated genome size that ranges from 375 to 625 Mb (Fig. 2a, Supplementary table 4). Notably, we found a *Parasponia*-specific expansion of ogre/tat LTR retrotransposons comprising 65 to 85 Mb (Fig. 2b). We then generated annotated reference genomes using high-coverage (~125X) sequencing of *P. andersonii* (accession WU1)^27^ and *T. orientalis* (accession RG33). These species were selected based on their low heterozygosity levels in combination with relatively small genomes. *T. tomentosa* was not used for a high-quality genome assembly because it is an allotetraploid (Supplementary Fig. 5, Supplementary Table 5-6).

**Figure 2:**
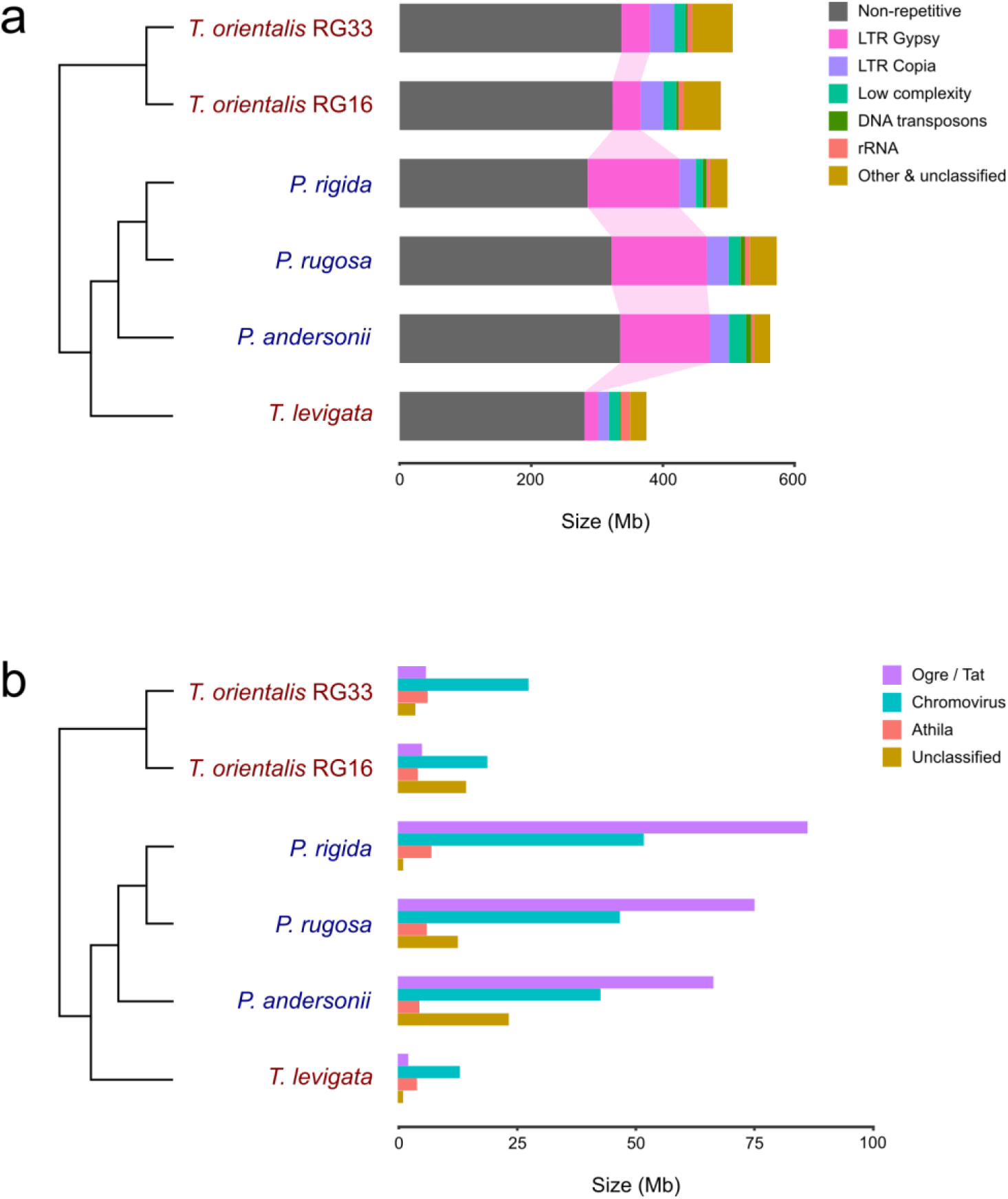
*Parasponia* and *Trema* genome structure. Estimated genome sizes and fractions of different classes of repeats as detected by RepeatExplorer, calibrated using k-mer based genome size estimates. **(a)** Total genome sizes and fractions of major repeat classes showing 1) a conserved size of around 300 Mb of non-repetitive sequence, and 2) a large expansion of gypsy-type LTR retrotransposons in all *Parasponia* compared with all *Trema* species. **(b)** Estimated size of gypsy-type LTR subclasses in *Parasponia* and *Trema* showing that expansion of this class was mainly due to a tenfold increase of Ogre/Tat to around 75Mb in *Parasponia*.

We generated orthogroups for *P. andersonii* and *T. orientalis* genes and six other Eurosid species, including arabidopsis (*Arabidopsis thaliana*) and the legumes medicago and soybean. From both *P. andersonii* and *T. orientalis* ~35,000 genes could be clustered into >29,000 orthogroups (Supplementary Table 7-8). Within these orthogroups we identified 25,605 *P. andersonii* - *T. orientalis* orthologous gene pairs based on phylogenetic analysis as well as whole genome alignments (Supplementary Table 8, note that there can be multiple orthologous gene pairs per orthogroup). These orthologous gene pairs had a median percentage nucleotide identity of 97% for coding regions (Supplementary Fig. 9–10). This further supports the recent divergence of the two species and facilitates their genomic comparison.

### Common utilization of symbiosis genes in *Parasponia* and medicago

To assess commonalities in the utilization of symbiosis genes in *Parasponia* species and legumes we employed two strategies. First, we identified close homologs of genes that were characterized to function in legume-rhizobium symbiosis. This revealed that *P. andersonii* contains orthologs of the vast majority of these legume symbiosis genes (117 out of 124) (Supplementary Table 1, Supplementary Data File 1). Second, we compared the sets of genes with enhanced expression in nodules of *Parasponia* and medicago. RNA sequencing of *P. andersonii* nodules revealed 1,725 genes that have a significantly enhanced expression level (fold change > 2, p < 0.05, DESeq2 Wald test) in any of three nodule developmental stages compared with uninoculated roots (Supplementary Fig. 11; Supplementary Table 9). For medicago, we used a comparable set of nodule-enhanced genes (1,463 genes)^31^. We then determined the overlap of these two gene sets based on orthogroup membership and found that 102 orthogroups comprise both *P. andersonii* and medicago nodule-enhanced genes. This number is significantly larger than is to be expected by chance (permutation test, p < 0.02)(Supplementary Fig. 12). Based on phylogenetic analysis of these orthogroups we found that in 85 cases (out of 1,725) putative orthologs have been utilized in both *P. andersonii* and medicago root nodules (Supplementary Table 10, Supplementary Data File 2). Among these 85 commonly utilized genes are 15 that we have identified in the first strategy; e.g. the LCO-responsive transcription factor *NIN* and its downstream target *NUCLEAR TRANSCRIPTION FACTOR*-*YA1* (*NFYA1*) that are essential for nodule organogenesis^16,17,32,33^, and *RHIZOBIUM DIRECTED POLAR GROWTH* (*RPG*) involved in intracellular infection^34^. A notable exception to this pattern of common utilization are the oxygen-binding hemoglobins. Earlier studies showed that *Parasponia* and legumes have recruited hemoglobin genes by divergent evolution^35^. Whereas legumes use class II LEGHEMOGLOBIN to control oxygen homeostasis, *Parasponia* recruited the paralogous class I HEMOGLOBIN 1 (HB1) for this function (Fig. 3a,b).

**Figure 3:**
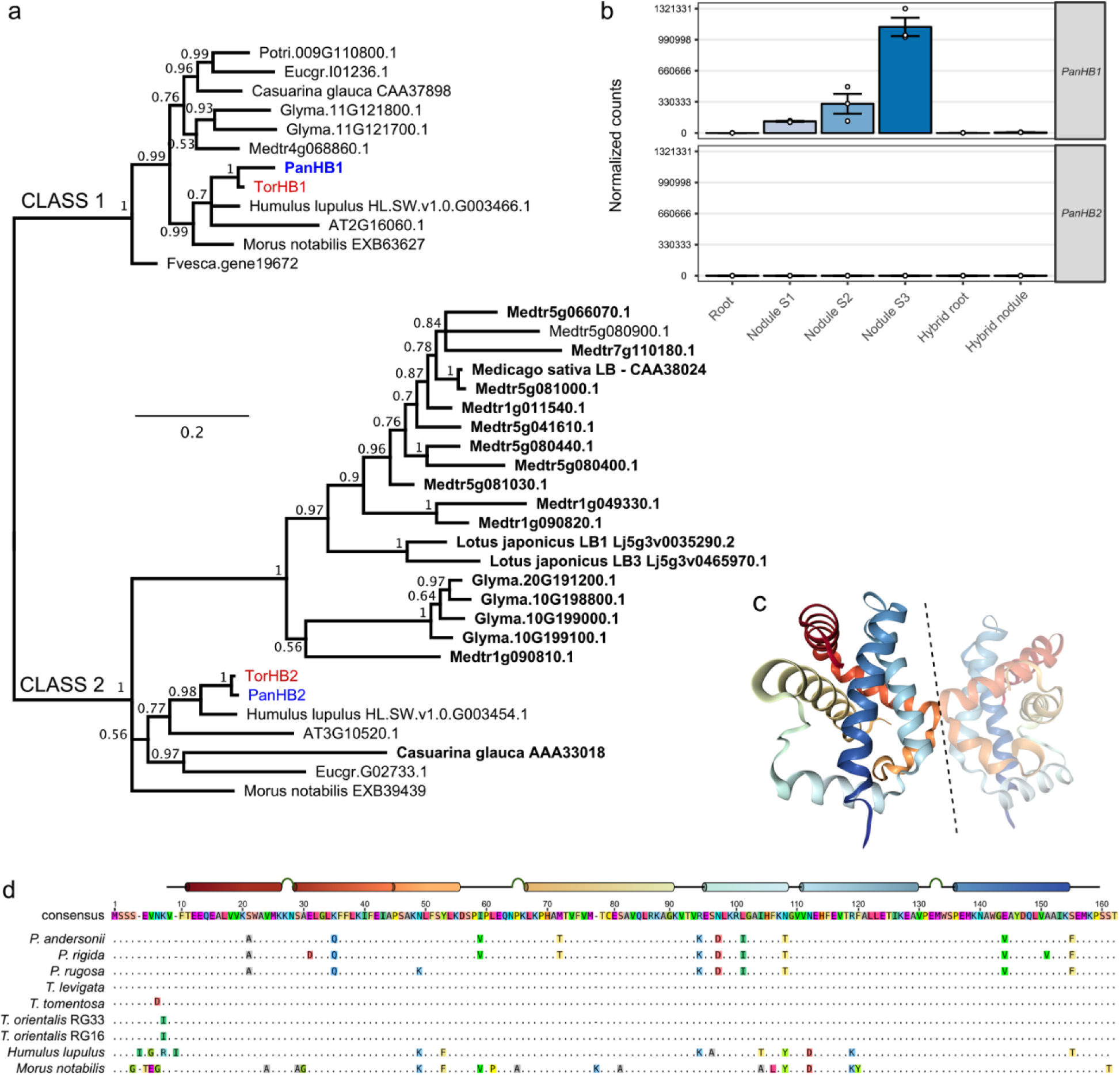
*Parasponia*-specific adaptations in class 1 hemoglobin protein HB1. **(a)** Phylogenetic reconstruction of class 1 (OG0010523) and class 2 hemoglobins (OG0002188). Symbiotic hemoglobins are marked in bold; legumes and the actinorhizal plant *casuarina* have recruited class 2 hemoglobins for balancing oxygen levels in their nodules. Conversely, *Parasponia* has recruited a class 1 hemoglobin *HB1* confirming parallel evolution of symbiotic oxygen transport in this lineage. *Medicago truncatula* (Medtr); *Glycine max* (Glyma), *Populus trichocarpa* (Potri); *Fragaria vesca* (Fvesca); *Eucalyptus grandis* (Eugr); *Arabidopsis thaliana* (AT). Node values indicate posterior probabilities; Scale bar represents substitutions per site. *Parasponia* marked in blue, *Trema* in red. **(b)** Expression profile of *PanHB1* and *PanHB2* in *P. andersonii* roots, stage 1-3 nodules, and in *P. andersonii* × *T. tomentosa* F_1_ hybrid roots and nodules (line H9). Expression is given in DESeq2 normalized read counts, error bars represent standard error of three biological replicates, dots represent individual expression levels. **(c)** Crystal structure of the asymmetric dimer of PanHB1 as deduced by Kakar *et al*. 2011^36^. Dashed line separates the two units. **(d)** Protein sequence alignment of class 1 hemoglobins from *Parasponia* spp., *Trema* spp., *Humulus lupulus*, and *Morus notabilis*. Only amino acids that differ from the consensus are drawn. A linear model of the crystal structure showing alpha helices and turns is depicted above the consensus sequence. There are seven amino acids that consistently differ between all *Parasponia* and all *Trema* species we sampled: Ala(21), Gln(35), Asp(97), Ile(101), Thr(108), Val(144), and Phe(155). These differences therefore correlate with the functional divergence between *P. andersonii* PanHB1 and *T. tomentosa TtoHB1^35,36^*. All seven consistently different sites are identical for all sampled *Trema* species, and five are identical for *Trema*, *Humulus*, and *Morus;* at both the amino acid and nucleotide level. This shows that these sites are conserved in all species except *Parasponia* and therefore supports adaptation of HB1 in the common ancestor of *Parasponia*. This suggests that the ancestral form of HB1 had oxygen affinities and kinetics that were not adapted to rhizobium symbiosis.

By exploiting the insight that nodule organogenesis and rhizobial infection can be genetically dissected using hybrid plants we classified these commonly utilized genes into two categories based on their expression profiles in both *P. andersonii* and F1 hybrid roots and nodules (Fig. 4). The first category comprises genes that are upregulated in both *P. andersonii* and hybrid nodules and that we associate with nodule organogenesis. The second category comprises genes that are only upregulated in the *P. andersonii* nodule that we therefore associate with infection and/or fixation. These variations in expression show that the commonly utilized genes commit functions in various developmental stages of the *P. andersonii* root nodule.

**Figure 4:**
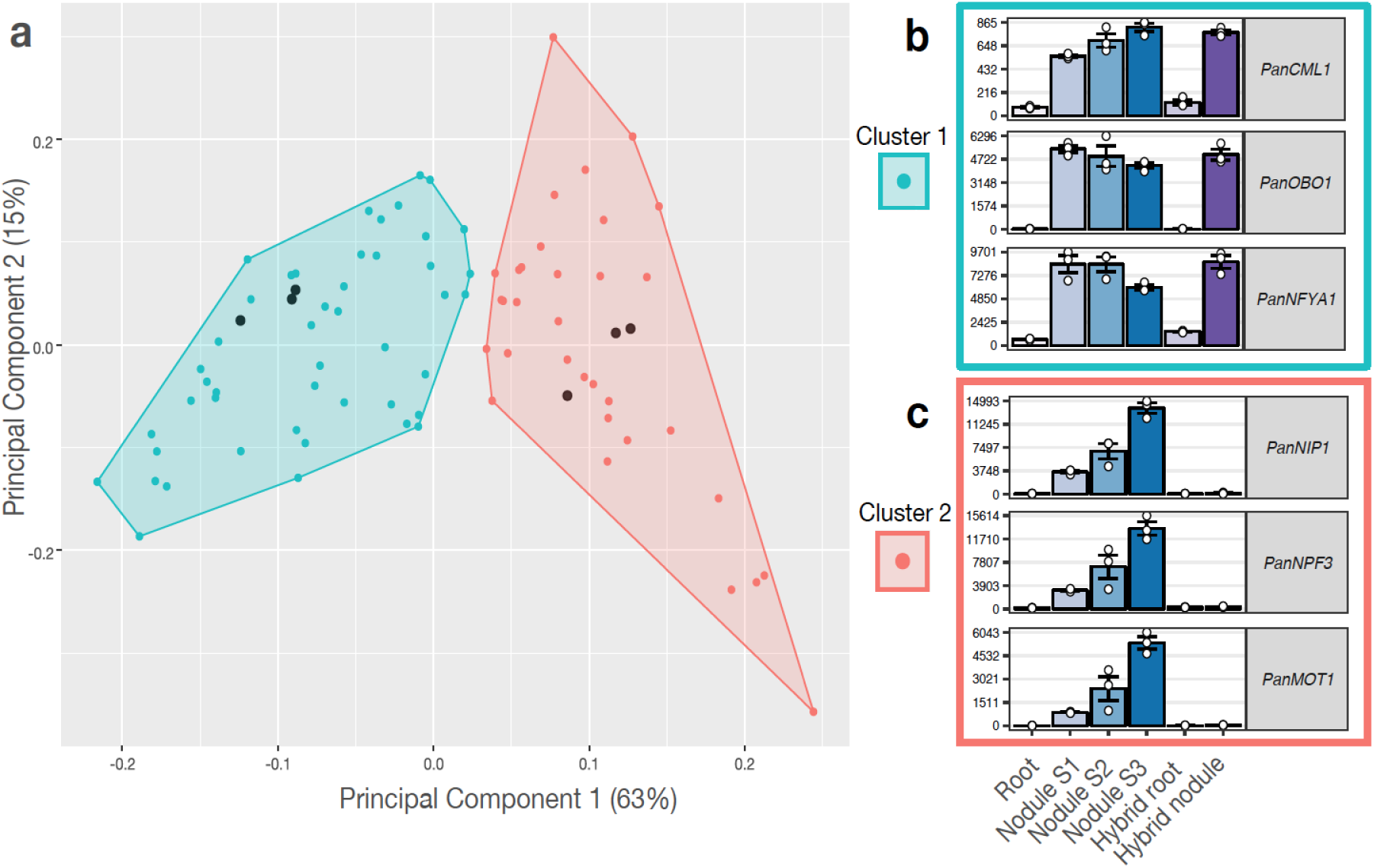
Clustering of commonly utilized symbiosis genes based on expression profile. **(a)** Principal component analysis plot of the expression profile of 85 commonly utilized symbiosis genes in 18 transcriptome samples: *P. andersonii* roots and nodules (stage 1-3), hybrid roots and nodules (line H9). All samples have three biological replicates. First two components are shown, representing 78% of the variation in all samples. Colors indicate clusters (K-means clustering using pearson correlation as distance measure, k=2) of genes with similar expression patterns. The three genes with the highest pearson correlation to the cluster centroids are indicated as black dots. **(b-c)** Expression profiles of representative genes for each cluster. **(b)** Cluster 1 represents genes related to organogenesis: these genes are upregulated in both *P. andersonii* and hybrid nodules. **(c)** Cluster 2 represents genes related to infection and fixation: these genes are highly upregulated in *P. andersonii* nodules, but do not respond in the hybrid nodule. *PanCML1: P. andersonii CALMODULIN 1; PanOBOl: P. andersonii ORGAN BOUNDARY*-*LIKE 1; PanNFYAI: P. andersonii NUCLEAR TRANSCRIPTION FACTOR*-*YA 1; PanNIP: P. andersonii AQUAPORIN NIP NODULIN26-LIKE; PanNPF3: P. andersonii NITRATE/PEPTIDE TRANSPORTER FAMILY 3; PanMOTI: P. andersonii MOLYBDATE TRANSPORTER 1*.

### Lineage-specific adaptation in *Parasponia* HEMOGLOBIN 1

We further examined *HB1* as it was recruited independently from legumes (Fig. 3a,b)^35^. Biochemical studies have revealed that *P. andersonii* PanHB1 has oxygen affinities and kinetics that are adapted to their symbiotic function, whereas this is not the case for *T. tomentosa TtoHBI^35,36^*. We therefore examined HB1 from *Parasponia* species, *Trema* species, and other non-symbiotic Rosales species to see if these differences are due to a gain of function in *Parasponia* or a loss of function in the non-symbiotic species. Based on protein alignment we identified *Parasponia*-specific adaptations in 7 amino acids (Fig. 3c,d). Among these is Ile(101) for which it is speculated to be causal for a functional change in *P. andersonii* HB1^36^. HEMOGLOBIN-controlled oxygen homeostasis in rhizobium-infected nodule cells is crucial to protect the nitrogen-fixing enzyme complex Nitrogenase^2,3^. Therefore, *Parasponia*-specific gain of function adaptations in *HB1* most likely were an essential evolutionary step towards functional rhizobium nitrogen fixing root nodules.

### Parallel loss of symbiosis genes in *Trema* and other relatives of *Parasponia*

Evolution of complex genetic traits is often associated with gene copy number variations (CNVs)^37^. To test if CNVs were associated with a potential independent evolution of nodulation in *Parasponia*, we focussed on two gene sets: (1) the 117 symbiosis genes that have been characterized in legumes, and (2) the 1,725 genes with a nodule-enhanced expression in *P. andersonii* (these sets partially overlap and add up to 1,791 genes; see Supplementary Fig. 13). To ensure that our findings are consistent between the *Parasponia* and *Trema* genera and not due to species-specific events, we analyzed the additional draft genome assemblies of two *Parasponia* and two *Trema* species (Supplementary Table 6). Finally, we discarded *Trema*-specific duplications as we considered them irrelevant for the nodulation phenotype. This resulted in only 11 consistent CNVs in the 1,791 symbiosis genes examined, further supporting the recent divergence between *Parasponia* and *Trema*. Due to the dominant inheritance of nodule organogenesis in F_1_ hybrid plants, we anticipated finding *Parasponia*-specific gene duplications that could be uniquely associated with nodulation. Surprisingly, we found only one consistent *Parasponia*-specific duplication in symbiosis genes; namely for a *HYDROXYCINNAMOYL*-*COA SHIKIMATE TRANSFERASE* (*HCT*) (Supplementary Fig. 14–15). This gene has been investigated in the legume forage crop alfalfa (*Medicago sativa*), where it was shown that HCT expression correlates negatively with nodule organogenesis^38,39^. Therefore, we do not consider this duplication relevant for the nodulation capacity of *Parasponia*. Additionally, we identified three consistent gene losses in *Parasponia* among which is the ortholog of LysM-type *EXOPOLYSACCHARIDE RECEPTOR 3* that in lotus inhibits infection of rhizobia with incompatible exopolysaccharides^40,41^ (Table 1, Supplementary Fig. 16–17). Such gene losses may have contributed to effective rhizobium infection in *Parasponia* and their presence in *T. tomentosa* could explain the lack of intracellular infection in the F1 hybrid nodules.

**Table 1.** Copy number variants in nodulation genes that are consistent between *Parasponia* and *Trema* genera.

| Name | ID | CNV type | Class | Description |
| --- | --- | --- | --- | --- |
| **PanNFP2** | PanWU01x14_asm01_ann01_320250 | loss in *Trema* | LS,NE | LysM domain containing receptor kinase, putative rhizobium LCO receptor |
| **PanCRK11** | PanWU01x14_asm01_ann01_285030 | loss in *Trema* | NE | Cysteine rich receptor like kinase |
| **PanLEK1** | PanWU01x14_asm01_ann01_069780 | loss in *Trema* | NE | Concanavalin A-like lectin receptor kinase |
| **PanNIN** | PanWU01x14_asm01_ann01_111140 | loss in *Trema* | LS,CR | Ortholog of transcription factor NODULE INCEPTION |
| **PanRPG** | PanWU01x14_asm01_ann01_272380 | loss in *Trema* | LS,CR | Ortholog of long coiled-coil protein RHIZOBIUM-DIRECTED POLAR GROWTH |
| **PanDEF1** | PanWU01x14_asm01_ann01_187760 | loss in *Trema* | NE | Defensin-like protein |
| **PanGAT** | PanWU01x14_asm01_ann01_150960 | loss in *Trema* | NE | Gamma-aminobutyric acid (GABA) transporter |
| **PanHCT1** | PanWU01x14_asm01_ann01_046570 | duplication in *Parasponia* | NE | Hydroxycinnamoyl-CoA shikimate / Quinate hydroxycinnamoyl transferase |
| **TorEPR** | TorRG33x02_asm01_ann01_052550 | loss in *Parasponia* | LS | LysM domain containing receptor kinase, putative rhizobium exopolysaccharide receptor |
| **TorN19L3** | TorRG33x02_asm01_ann01_066920 | loss in *Parasponia* | LS | NODULIN19-like protein |
| **TorIPT4** | TorRG33x02_asm01_ann01_307000 | loss in *Parasponia* | LS | Isopentenyltransferase |

Contrary to our initial expectations, we discovered consistent loss or pseudogenization of seven symbiosis genes in *Trema*. These genes have a nodule-specific expression profile in *Parasponia*, suggesting that they function exclusively in symbiosis (Fig. 5). Three of these are orthologs of genes that are essential for establishment of nitrogen-fixing nodules in legumes: *NIN*, *RPG*, and the LysM-type LCO receptor *NFP*/*NFR5* (Supplementary Fig. 18–19). In the case of *NFP/NFR5*, we found two close homologs of this gene, *NFP1* and *NFP2*, of which the latter is consistently pseudogenized in *Trema* species (Fig. 6). In an earlier study we used RNA interference (RNAi) to target *PanNFP1*, which led to reduced nodule numbers and a block of intracellular infection by rhizobia as well as arbuscular mycorrhiza^27^. Most probably, however, the RNAi construct unintentionally also targeted *PanNFP2*, as both genes are 69% identical in the 422 bp RNAi target region. Phylogenetic reconstruction revealed that the *NFP1*-*NFP2* duplication predates the divergence of legumes and *Parasponia*, and that *Parasponia NFP2* is most closely related to legume *MtNFP/LjNFR5* rhizobium LCO receptors (Fig. 6). Additionally, in *P. andersonii* nodules *PanNFP2* is significantly higher expressed than *PanNFP1* (Supplementary Fig. 20). Taken-together, this suggests that *PanNFP2* represents a rhizobium LCO receptor that functions in nodule formation and intracellular infection in *Parasponia*.

**Figure 5:**
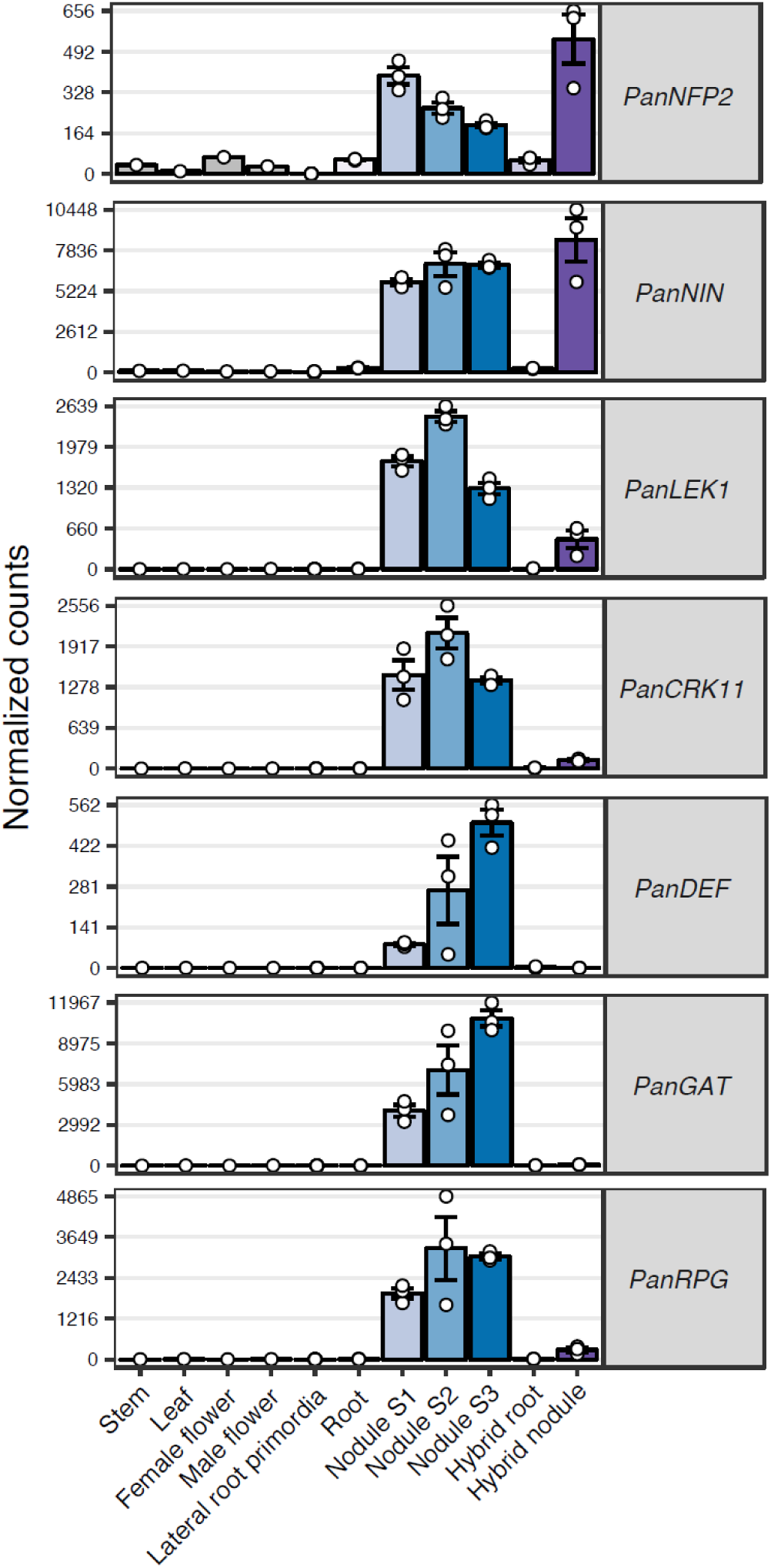
Expression profile of *Parasponia* symbiosis genes that are lost in *Trema* species. Expression of symbiosis genes in *P. andersonii* stem, leaf, female and male flowers, lateral root primordia, roots and 3 nodule stages (S1-3), and in *P. andersonii × T. tomentosa* F_1_ hybrid roots and nodules (line H9). Expression is given in DESeq2 normalized read counts, error bars represent standard error of three biological replicates for lateral root primordia, root, and nodule samples. Dots represent individual expression levels. *PanNFP2: P. andersonii NOD FACTOR PERCEPTION 2*, *PanNIN: P. andersonii NODULE INCEPTION*, *PanLEKI: P. andersonii LECTIN RECEPTOR KINASE 1*, *PanCRK11: P. andersonii CYSTEINE*-*RICH RECEPTOR KINASE 11*, *PanDEFI: P. andersonii DEFENSIN 1; PanRPG: P. andersonii RHIZOBIUM DIRECTED POLAR GROWTH*.

**Figure 6:**
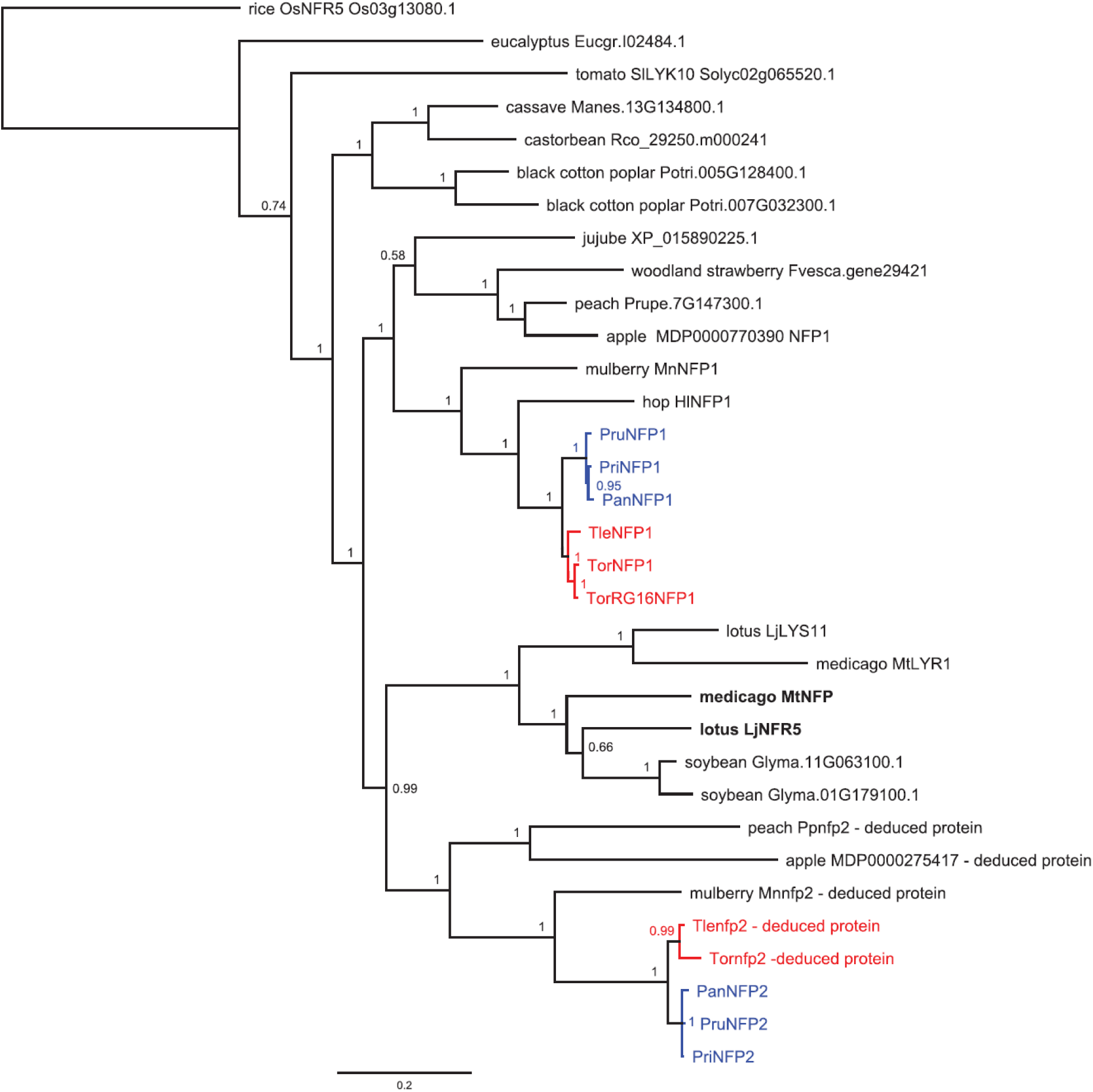
*Parasponia* NFP1 and NFP2 are homologous to legume LCO receptors MtNFP/LjNFR5. Phylogenetic reconstruction of the NFP/NFR5 orthogroup. NFP2 protein sequences of *T. levigata*, *T. orientals*, *Moms notabilis*, *Malus* × *domestica* and *Prunus persica* have been deduced from pseudogenes. Included species: *Parasponia andersonii* (Pan); *Parasponia rigida* (Pri); *Parasponia rugosa* (Pru); *Trema orientalis* RG33 (Tor); *Trema orientalis* RG16 (TorRG16); *Trema levigata* (Tle); medicago (*Medicago truncatula*, Mt); lotus (*Lotus japonicus*, Lj); soybean (*Glycine max*, Glyma); peach (*Prunus persica*, Ppe); woodland strawberry (*Fragaria vesca*, Fvesca); black cotton poplar (*Populus trichocarpa*, Potri); eucalyptus (*Eucalyptus grandis*, Eugr); jujube (*Ziziphus jujube*), apple (*Malus × domestica*), mulberry (*Morus notabilis)*, hops (*Humulus lupulus* (natsume.shinsuwase.v1.0)), cassave (*Manihot esculenta*), rice (*Oryza sativa*), tomato (*Solanum lycopersicum*), castor bean (*Ricinus communis*). Node numbers indicate posterior probabilities, scale bar represents substitutions per site. *Parasponia* proteins are marked in blue, *Trema* in red.

Based on expression profiles and phylogenetic relationships we postulate that also *Parasponia NIN* and *RPG* commit essential symbiotic functions similar as in other nodulating species (Fig. 5; Supplementary Fig. 21–23). Expression of *PanRPG* increases >300 fold in *P. andersonii* nodules that become intracellularly infected (nodule stage 2), whereas in F_1_ hybrid nodules-which are devoid of intracellular rhizobium infection-*PanRPG* upregulation is less than 20-fold (Fig. 5). This suggests that *PanRPG* commits a function in rhizobium infection, similar as found in medicago^34^. The transcription factor *NIN* has been studied in several legume species as well as in the actinorhizal plant casuarina (*Casuarina glauca*) and in all cases shown to be essential for nodule organogensis^14,16,42,43^. Loss of *NIN* and/or *NFP2* in *Trema* species can explain the genetic dominance of nodule organogenesis in the *Parasponia* × *Trema* F1 hybrid plants.

Next, we questioned whether loss of these symbiosis genes also occurred in more distant relatives of *Parasponia*. We analysed non-nodulating species representing 6 additional lineages of the Rosales clade; namely hops (*Humulus lupulus*, Cannabaceae)^44^, mulberry (*Morus notabilis*, Moraceae)^45^, jujube (*Ziziphus jujuba*, Rhamnaceae)^46^, peach (*Prunus persica*, Rosaceae)^47^, woodland strawberry (*Fragaria vesca*, Rosaceae)^48^, and apple (*Malus × domestica*, Rosaceae)^49^. This revealed a consistent pattern of pseudogenization or loss of *NFP2*, *NIN* and *RPG* orthologs, the intact jujube *ZjNIN* being the only exception (Fig. 7). We note that for peach *NIN* was previously annotated as protein-coding gene^47^. However, based on comparative analysis of conserved exon structures we found two out-of-frame mutations (see supplementary Fig. 24). Because the pseudogenized symbiosis genes are largely intact in most of these species and differ in their deleterious mutations, the loss of function of these essential symbiosis genes should have occurred recently and in parallel in at least seven Rosales lineages. As we hypothesize that *NFP2*, *NIN* and *RPG* are essential for nodulation, we argue that *Trema* species, hops, mulberry, jujube, woodland strawberry, apple, and peach irreversibly, recently, and independently lost the potential to nodulate.

**Figure 7:**
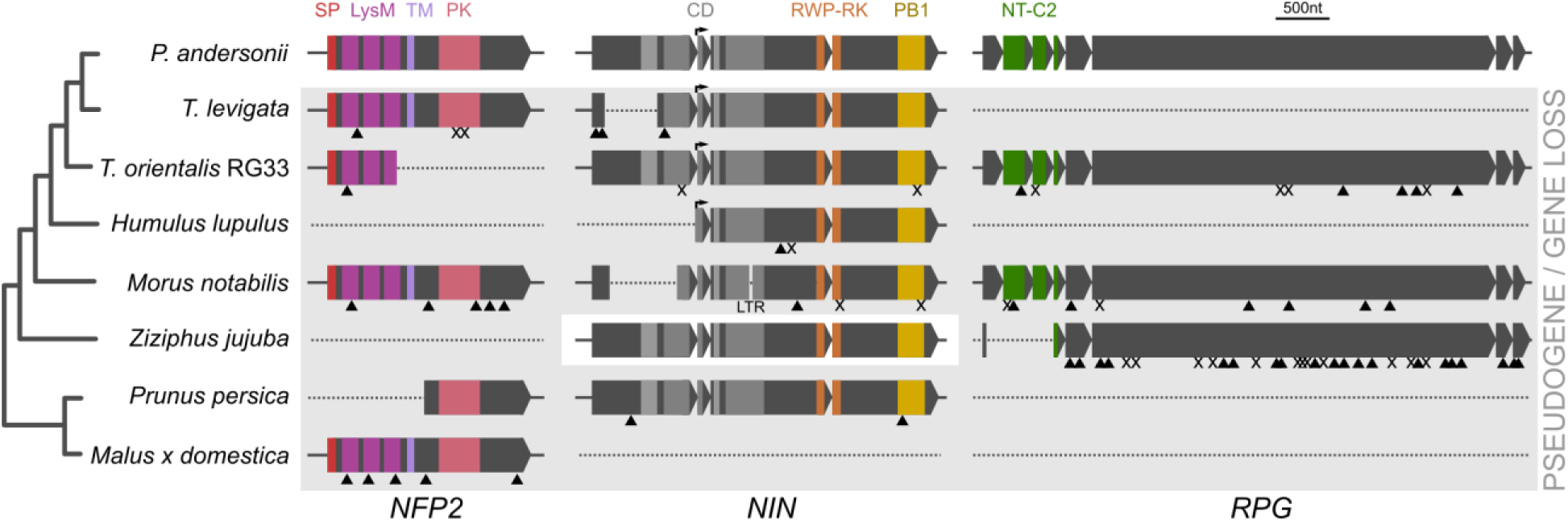
Parallel loss of symbiosis genes in non-nodulating Rosales species. Pseudogenization or loss of *NOD FACTOR PERCEPTION 2* (*NFP2*), *NODULE INCEPTION* (*NIN*) and *RHIZOBIUM*-*DIRECTED POLAR GROWTH (RPG)* in two phylogenetically independent *Trema* lineages, *Humulus lupulus*, *Morus notabilis*, *Prunus persica*, and *Malus × domestica*. In *Ziziphus jujuba NFP2* is lost and *RPG* is pseudogenized, but *NIN* is intact. In *Fragaria vesca* all three genes are lost (not shown). Introns are indicated but not scaled. Triangles indicate frame-shifts; X indicate premature stop codons; LTR indicates long terminal repeat retrotransposon insertion (not scaled); arrows indicate alternative transcriptional start site in *NIN*. SP = signal peptide (red); LysM: 3 Lysin Motif domains (magenta); TM = transmembrane domain (lilac); PK = protein kinase (pink); CD = 4 conserved domains (grey); RWP-RK: conserved amino acid domain (orange); PB1 = Phox and Bem1 domain (yellow); NT-C2 = N-terminal C2 domain (green).

## DISCUSSION

Here we present the nodulating Cannabaceae species *Parasponia* as a comparative system to obtain insights in the evolutionary trajectory of nitrogen-fixing symbioses. Instead of finding gene duplications that can explain a gain of symbiosis in *Parasponia*, we found parallel loss or pseudogenization of symbiosis genes in non-nodulating Rosales species. This indicates that in non-nodulating Rosales lineages these symbiosis genes experienced a recent period of reduced functional constraints. This challenges current hypotheses on the evolution of nitrogen fixing plant-microbe symbiosis.

Evolution of nodules is widely considered to be a two-step process: first an unspecified predisposition event in the ancestor of all nodulating species, bringing species in the nitrogen fixation clade to a precursor state for nodulation^1,6^. Subsequently, nodulation originated in parallel; eight times with *Frankia* and twice with rhizobium^1,6–9^. This hypothesis is most parsimonious and suggests a minimum number of independent losses of symbiosis. *NFP/NFR5*, *NIN* and *RPG* are essential for nodulation in legumes and-in case of *NIN*-the non-legume casuarina^43^. Consequently, the non-nodulating species that have lost these genes irreversibly lost the potential to nodulate. This opposes the current view that non-host relatives of nodulating species are generally in a precursor state for nodulation^1,6^.

The loss of symbiosis genes in non-nodulating plants is difficult to explain under the current hypothesis of parallel origins of nodulation. These genes commit functions that currently cannot be linked to any non-symbiotic processes. As a consequence, the interpretation of why such a diverse set of genes repeatedly experienced reduced functional constraints in non-nodulating plant lineages requires these genes to be linked to some other, yet unknown, common process. Additionally, the current hypothesis of parallel origins would imply convergent recruitment of at least 85 genes to commit symbiotic functions in *Parasponia* and legumes. This implies parallel evolution of a highly complex trait.

Alternatively, the parallel loss of symbiosis genes in non-nodulating plants can be interpreted as a single gain and massive loss of nodulation. In this new hypothesis nodulation is much older than generally anticipated, and possibly represents the hypothetical predisposition event in the nitrogen fixation clade. Subsequently, nodulation was lost in most descendent lineages (hence massive). This new hypothesis fits our data better in four ways. (I.) It more convincingly explains the parallel loss of symbiosis genes in non-nodulating plants, because then gene loss correlates directly with loss of nodulation. (II.) A single gain of nodulation explains the origin of the conserved set of more than 80 symbiosis genes utilized by *Parasponia* and medicago. (III.) It is in line with the lack of *Parasponia*-specific gene duplications that associate with nodulation. (IV.) The duplication in the NFP/NFR5 clade encoding putative LCO LysM-type receptors predates the Rosales and Fabales split, thereby coinciding with the origin of the nitrogen fixation clade. A single gain of nodulation would require only a single (sub)neofunctionalization event of LCO-receptors to function in root nodule formation. Additionally, the single gain-massive loss hypothesis eliminates the predisposition event, a theoretical concept that currently cannot be addressed mechanistically. Therefore, we consider a single gain-massive loss hypothesis as plausible.

Loss of nodulation is not controversial, as it is generally considered to have occurred >20 times in the legume family^6,7^. Nevertheless, the single gain-massive loss hypothesis implies many more events than the current hypothesis of parallel gains. Based on phylogenetic evidence, the minimum number of losses required to explain the pattern of nodulation in the nitrogen fixation clade implies 20 events in Rosales (8 of which in the Cannabaceae), 5 in Fagales, 3 in Cucurbitales and 2 in Fabales (not taken into account Fabaceae)^7^. However, as the identified pseudogenes in *Trema* species, mulberry, jujube, apple, and peach are relatively intact we hypothesize that loss of nodulation has occurred relatively recent, which would imply significantly more events. In either case, this hypothesis is not the most parsimonious. On the other hand, it is conceptually easier to lose a complex trait, such as nodulation, rather than to gain it^9^. Genetic studies in legumes indeed demonstrated that nitrogen-fixing symbiosis can be abolished by a single knockout mutation in tens of different genes, among which are *NFP/NFR5*, *NIN* and *RPG* (Supplementary Table 1). This suggests that simple parsimony may not be the best way to model the evolution of nodulation.

Massive, recent and parallel loss of nodulation may have been triggered by changes at a geological scale, e.g. a glacial maximum. During periods of glacial maxima, which occurred between 18,000 and 800,000 years ago, atmospheric CO_2_ levels dropped below 200 ppm^50,51^. Experiments show that such CO_2_ concentrations have profound effects on photosynthesis and plant growth in general^52^. Under such conditions photosynthates may have been the growth limiting factor, rather than fixed nitrogen^52–54^. In line with this nitrogen-fixation rates in the legume *Prosopis glandulosa* (honey mesquite) can drop to zero when grown at 200 ppm CO_2_^53^ Therefore it is likely that the nitrogen fixation trait has experienced relaxed constraints during periods of low atmospheric CO_2_ concentration, leading to genetic defects.

Based on the single gain-massive loss hypothesis we can make the following predictions. First, the hypothesis implies that many (if not most) ancestral species in the nitrogen-fixing clade were nodulators. This should be substantiated by fossil evidence. Currently, fossil data on nodules are basically absent with only a single report on a fossilized nodule that is estimated to be 11,5 thousand years old^55,56^. An alternative strategy is to infer the presence of nitrogen-fixing symbiosis from N isotope variation in fossil tree rings. This method was successfully applied to discriminate tree species that predominantly utilize biologically fixed nitrogen from tree species that use nitrogen resources retrieved from soil^57^. Secondly, we predict that actinorhizal plant species maintained *NIN*, *RPG*, and possibly *NFP2* (in case LCOs are used as symbiotic signal^58^), and that these genes are essential for nodulation. This can be shown experimentally, as was done for *NIN* in casuarina^43^.

The loss of symbiosis genes in non-nodulating plant species is not absolute, as we observed a functional copy of *NIN* in jujube. This pattern is similar to the pattern of gene loss in species that lost endomycorrhizal symbiosis^59,60^. Also in that case, occasionally such genes have been maintained in non-mycorrhizal plants. Conservation of *NIN* in jujube suggests that this gene has a non-symbiotic function. Contrary to *NFP2*, which is the result of a gene duplication near the origin of the nitrogen-fixing clade, functional copies of *NIN* are also present in species outside the nitrogen-fixing clade (Supplementary Fig. 22). This suggests that these genes may have retained-at least in part-an unknown ancestral non-symbiotic function in some lineages within the nitrogen-fixing clade. Alternatively, *NIN* may have acquired a new non-symbiotic function within some lineages in the nitrogen-fixing clade.

As hemoglobin is crucial for rhizobium symbiosis^3^, it is striking that *Parasponia* and legumes do not use orthologous copies of hemoglobin genes in their nodules. At first sight this seems inconsistent with a single gain of nodulation. However, a scenario that incorporates a switch in microsymbionts can reconcile the use of paralogous hemoglobin genes with the occurrence of two types of microsymbiont in the nitrogen fixing clade. This scenario would dictate a single gain of actinorhizal symbiosis in the nitrogen fixing clade, and a switch from *Frankia* to rhizobium in the ancestors of both *Parasponia* and legumes. As *Frankia* species possess intrinsic physical characteristics to protect the Nitrogenase enzyme for oxidation, expression of plant encoded hemoglobin in nodules is not a prerequisite for nitrogen fixation in actinorhizal plants^61–64^. In line with this, there is no evidence that *Ceanothus* spp. (Rhamnaceae, Rosales) - which represent the closest nodulating relatives of *Parasponia* - express a hemoglobin gene in *Frankia*-infected nodules^62–64^. A microsymbiont switch from *Frankia* to rhizobium would therefore require adaptations in hemoglobin. Based on the fact that *Parasponia* acquired lineage-specific adaptations in HB1 that are considered to be essential to control oxygen homeostasis in rhizobium root nodules^35,36^, such a symbiont switch may have occurred early in the *Parasponia* lineage.

The uncovered evolutionary trajectory of a rhizobium nitrogen-fixing symbiosis provides novel leads in attempts to engineer nitrogen-fixing root nodules in agricultural crop plants. Such a translational approach is anticipated to be challenging^65^, and the only published study so far, describing transfer of 8 LCO signaling genes, was unsuccessful^66^. If we interpret the parallel loss of symbiosis genes in non-nodulating plants as evidence that these genes have been neofunctionalized to commit symbiotic functions, then this gene set is essential in any engineering approach. However, transfer of symbiosis genes may not be sufficient to obtain functional nodules. The lack of infection in nodules on hybrid plants that contain a full genome complement of *Parasponia* indicates the presence of an inhibitory mechanism in *T. tomentosa*. Such a mechanism may also be present in other non-host species. Consequently, engineering nitrogen-fixing nodules requires gene knockouts in non-nodulating plants to overcome inhibition of intracellular infection. *Trema* may be the best candidate species for such a (re)engineering approach, due to its high genetic similarity with *Parasponia* and the availability of transformation protocols^67^. Therefore, the presented *Parasponia*-*Trema* comparative system may not only be suited for evolutionary studies, but also can form an excellent experimental platform to obtain essential insights to engineer nitrogen fixing root nodules.

## MATERIAL AND METHODS

### *Parasponia* - *Trema* intergeneric crossing and hybrid genotyping

*Parasponia* and *Trema* are wind-pollinated species. A female-flowering *P. andersonii* individual WU1.14 was placed in a plastic shed together with a flowering *T. tomentosa* WU10 plant. Putative F_1_ hybrid seeds were germinated (see Supplementary Methods) and transferred to potting soil. To confirm the hybrid genotype a PCR marker was used that visualizes a length difference in the promoter region of *LIKE*-*AUXIN 1* (*LAX1*) (primers: LAX1-f: ACATGATAATTTGGGCATGCAACA, LAX1-r: TCCCGAATTTTCTACGAATTGAAA, amplicon size *P. andersonii:* 974 bp; *T. tomentosa:* 483 bp). Hybrid plant H9 was propagated *in vitro^27,68^*. The karyotype of the selected plants was determined according to Geurts and De Jong 2013^69^.

### Nodulation and nitrogenase activity assays

All nodulation assays were conducted with *Mesorhizobium plurifarium* BOR2. This strain was isolated from *P. andersonii* root nodules grown in soil samples collected from the root rhizosphere of *Trema orientalis* plants in Malaysian Borneo, province of Sabah^70^. *M. plurifarium* was grown on yeast extract mannitol medium at 28°C^30^. Plants were grown in sterile plastic 1 liter pots containing perlite and EKM medium supplemented with 0.375 mM NH_4_NO_3_ and rhizobium (OD600:0.05)^71^. Nodule number per plant was quantified 6 weeks post inoculation.

Acetylene reduction assays^72^ were conducted on nodules harvested 6 weeks post inoculation with *Mesorhizobium plurifarium* strain BOR2. Nodules were sampled per plant and collected in 15 ml headspace vials with screw lids. 2.5 ml of acetylene was injected into the vial and incubated for about 10 minutes, after which 1 ml headspace was used to quantify ethylene nitrogenase activity using an ETD 300 detector (Sensor Sense, Nijmegen, The Netherlands; Isogen, Wageningen, The Netherlands)^73^.

To isolate *P. andersonii* nodules at 3 developmental stages nodules were separated based on morphology and size. Stage 1: nodules are round and < 1mm in diameter in size. The outer cell layers of stage 1 nodules are transparent. Light microscopy confirmed that at this stage, rhizobia already reach the central part of the nodule, but are mainly present in the apoplast (Fig. 1h). Stage 2: nodules are brownish, and ~2 mm in size. Nodules have formed an apical meristem and 2-3 cell layers have been infected by rhizobia (Fig. 1i). Stage 3: nodules are pinkish on the outside due to accumulation of haemoglobin and > 2 mm in size. Light microscopy showed that stage 3 nodules contain zones of fully infected cells (Fig. 1j). For each of these stages three biological replicates were used for RNA sequencing.

### Arbuscular mycorrhization assay

Two week old seedlings were transferred to 800 ml Sand:Granule*:Rhizophagus irregularis* (*Rir*, INOQ TOP-INOQ GmbH, Schnega Germany) inoculum mixture (1:1:0.01), irrigated with 80 ml ½ strength modified Hoagland solution containing 20 μM K_2_HPO_4_^74^ and grown for an additional 6 weeks at 28°C, under a photoperiod of 16/8h (day/night). 50 ml additional nutrient solution was provided once a week. Mycorrhization efficiency was analysed as previously described^75^ for three aspects: 1) frequency of fungal colonization in 1 cm root segments; 2) average level of mycorrhization in all root fragments, and 3) arbuscular abundance in all root fragments (Supplementary Fig. 1). Arbuscules were WGA-Alexafluor 488-stained and imaged according to Huisman *et al* 2015^76^.

### DNA/RNA sequencing

Paired-end Illumina libraries (insert size 500bp, 100bp reads) were prepared for all accessions (Supplementary Table 5), mate-pair libraries (3Kb, 7Kb, and 10Kb) and overlapping fragment libraries (450bp insert size, 250bp reads) were prepared for the reference accessions (*P. andersonii* accession WU01 and *T. orientalis* accession RG33). Paired-end and mate-pair libraries were sequenced on an Illumina HiSeq2000, overlapping libraries were sequenced on an Illumina MiSeq. For the *P. andersonii* and *T. orientalis* reference genomes a total of 75Gb (~132x genome coverage) and 61Gb (~121x coverage) of data was produced respectively. The other accessions were sequenced at an average coverage of ~30X. See Supplementary Methods for further details on library preparation and sequencing. RNA samples from various tissues and nodulation stages were isolated from *P. andersonii* and *Trema orientalis* RG33 (Supplementary Methods, Supplementary Table 11). Library preparation and RNA sequencing was conducted by B.G.I. (Shenzhen, China).

### Estimation of heterozygosity levels and genome size

To assess levels of heterozygosity and genome size we performed *k*-mer analyses. Multiplicities of 21-mers were extracted from the reads using Jellyfish (version 2.2.0)^77^ and processed using custom R scripts. First, a multiplicity threshold was determined below which most *k*-mers are considered to represent sequencing errors and which were excluded from further analysis. In principle, errors occur randomly and this generates a high frequency peak at multiplicity 1 after which frequency decreases and subsequently increases due to a broad frequency peak around the mean genome coverage. The error multiplicity threshold was therefore set at the multiplicity with the lowest frequency between these two peaks. Next, we identified the peak multiplicity as the one with the highest frequency. Homozygous genome coverage was estimated by scaling the peak multiplicity proportional to the difference of its frequency with that of multiplicities one below and above. Heterozygous coverage was defined as half that of the homozygous coverage (Supplementary Fig. 5). Finally, genome size was calculated as the total number of error-free *k*-mers divided by the estimated homozygous genome coverage (Supplementary Table 4). These estimates are generally comparable to those based on FACS measurements^78^ (Supplementary Table 3) except for genomes that differ much from the reference used to calibrate the FACS results (*Medicago truncatula*, ~500Mb). This inconsistency is probably due to the non-linearity of the FACS measurements. We therefore consider the quantitative genome estimates based on k-mer analysis a more accurate estimation of genome size.

### Characterization of repetitive sequences

Repetitive sequences are inherently difficult to assemble. We therefore characterized and quantified repetitive element using the ab initio graph-based clustering approach implemented in RepeatExplorer^79^. Analyses were based on random subsamples of 20,000 paired-end reads and included a reclustering step where clusters with shared mate pairs are merged (threshold k=0.2). Repeat classification was based on the RepeatExplorer Viridiplantae dataset and on plant organellar sequences. Relative sizes of repetitive sequences in the genome were scaled by the genome size estimations based on *k*-mer analysis to generate absolute sizes in Mb (Fig. 2).

### Assembly of reference genomes

The raw sequencing data were preprocessed. First, adapters (standard and junction) were removed and reads were trimmed using fastq-mcf (version 1.04.676)^80^. Minimum remaining sequence length was set to 50 for HiSeq data and 230 for MiSeq data. Duplicates were removed using FastUniq (version 1.1)^80,81^. Chloroplast and mitochondrial genomes were assembled first with IOGA (version 1) using reference sets of plant chloroplast and mitochondrial genomes^82^. Chloroplast and mitochondrial reads were identified and separated from the nuclear reads by mapping to four organellar assemblies (*Parasponia andersonii*, *Trema orientalis*, *Morus indica*, *Malus × domestica*) using BWA (version 0.7.10)^83^. Finally, a contamination database was produced by BLASTing contigs from earlier in-house draft genome assemblies from *Parasponia andersonii* and *Trema orientalis* against NCBIs nt database. Hits outside the plant kingdom were extracted using a custom script and corresponding sequences were downloaded from GenBank and a database of plant viruses was added (http://www.dpvweb.net/seqs/allplantfasta.zip). Genomics reads were cleaned by mapping to this contamination database.

The preprocessed data were *de novo* assembled using ALLPATHS-LG (release 48961)^84^. Relevant parameters were PLOIDY=2 and GENOME_SIZE=600000000. The assemblies were performed on the Breed4Food High Performance Cluster from Wageningen UR (http://breed4food.com).

Remaining contamination in the ALLPATHS-LG assembly was identified by blasting the assembled contigs to their respective chloroplast and mitochondrial genomes, the NCBI nr and univec databases (Downloaded 29 oktober 2014) and by mapping back genomic reads of the HiSeq 500bp insert size library. Regions were removed if they matched all of the following criteria: (1) significant blast hits with more than 98% identity (for the nr database only blast results that were not plant-derived were selected); (2) read coverage lower than 2 or higher than 50 (average coverage for the HiSeq 500bp insert size library is ~30x); (3) number of properly paired reads lower than 2.

Resulting contigs were subsequently scaffolded with two rounds of SSPACE (v3.0)^85^, standard with the mate pair libraries. In order to use reads mapped with BWA (v0.7.10) the SSPACE utility sam_bam2tab.pl was used. We used the output of the second run of SSPACE scaffolding as the final assembly.

Validation of the final assemblies showed that 90-100% of the genomic reads mapped back to the assemblies (Supplementary Table 5), and 94-98% of CEGMA^86^ and BUSCO^87^ genes were detected (Supplementary Table 6).

### Annotation of reference genomes

Repetitive elements were identified following the standard Maker-P recipe (http://weatherby.genetics.utah.edu/MAKER/wiki/index.php/Repeat_Library_Construction-Advanced accessed october 2015) as described on the GMOD site: (1) RepeatModeler with Repeatscout v1.0.5, Recon v1.08, RepeatMasker version open4.0.5, using RepBase version 20140131^88^ and TandemRepeatFinder; (2) GenomeTools: LTRharvest and LTRdigest^89^; (3) MITEhunter with default parameters^90^. We created species-specific repeat libraries for both *P. andersonii* and *T. orientalis* separately and combined these into a single repeat library, filtering out sequences that are >98% similar. We masked both genomes using RepeatMasker with this shared repeat library.

To aid the structural annotation we used 11 *P. andersonii* and 6 *T. orientalis* RNA sequencing datasets (Supplementary Table 11). All RNA-seq samples were assembled *de novo* using genome-guided Trinity^91^, resulting in one combined transcriptome assembly per species. In addition all samples were mapped to their respective reference genomes using BWA and processed into putative transcripts using cufflinks^92^ and transdecoder^93^. This resulted in one annotation file (gff) per transcriptome sample per species. As protein homolog evidence, only Swiss-Prot^94^ entries filtered for plant proteins were used. This way we only included manually verified protein sequences and prevented the incorporation of erroneous predictions. Finally, four gene-predictor tracks were used: 1) SNAP^95^, trained on *P. andersonii* transdecoder transcript annotations; 2) SNAP, trained on *T. orientalis* transdecoder transcript annotations; 3) Augustus^96^ as used in the BRAKER pipeline, trained on RNA-seq alignments^97^; 4) GeneMark-ET as used in the BRAKER pipeline, trained on RNA-seq alignments^98^.

First, all evidence tracks were processed by Maker-P^87,99^. The results were refined with EVidenceModeler (EVM)^100^, which was used with all the same tracks as Maker-P, except for the Maker-P blast tracks and with the addition of the Maker-P consensus track as additional evidence. Ultimately, EVM gene models were preferred over Maker-P gene models, except when there was no overlapping EVM gene model. Where possible, evidence of both species was used to annotate each genome (i.e. *de novo* RNA-seq assemblies of both species were aligned to both genomes).

To take maximum advantage of annotating two highly similar genomes simultaneously we developed a custom reconciliation procedure involving whole genome alignments. The consensus annotations from merging the EVM and Maker-P annotations were transferred to their respective partner genome using nucmer^101^ and RATT revision 18^102^ (i.e. the *P. andersonii* annotation was transferred to *T. orientalis* and *vice versa*), based on nucmer whole genome alignments (Supplementary Fig. 9). Through this reciprocal transfer, both genomes had two candidate annotation tracks, the original (called P and T) and the transferred (called P’ on *T. orientalis* and T’ on *P. andersonii*). This allowed for validation of both annotations simultaneously, assuming that two orthologous regions containing a single gene that has not changed since its common ancestor, should be annotated identically. If annotations between orthologous regions differ, we used RNA-seq evidence and protein alignments for curation, by picking one of four annotation combinations: P and T, P and T’, P’ and T, or P’ & T’. Picking one of these options is based on transcriptome coverage: the combination with the highest percentage of covered introns per annotation is the most likely. If there is insufficient coverage in any of the genomes, the combination with the highest pairwise identity based on protein alignments of the translated annotation is selected.

For the reconciliation procedure we developed a custom Python script. To deal with orthologous regions containing different numbers of annotations, we identified ‘annotation clusters’. This was done iteratively by selecting overlapping gene models and transferred gene models with the same gene ID. Two annotations were considered to be overlapping if they were on the same strand and at least one of each of their exons overlapped. This allowed for separate processing of genes on opposing strands, and ‘genes within genes’, i.e. a gene within the intron of another gene. The validation of annotation differences between *P. andersonii* and *T. orientalis* greatly reduces technical variation and improves all downstream analyses.

After automatic annotation and reconciliation 1,693 *P. andersonii* genes and 1,788 *T. orientalis* genes were manually curated. These were mainly homologs of legume symbiosis genes and genes that were selected based on initial data exploration.

To assign putative product names to the predicted genes we combined BLAST results against Swiss-Prot, TrEMBL and nr with InterProScan results (custom script). To annotate GO terms and KEGG enzyme codes Blast2GO was used with the nr BLAST results and interproscan results. Finally, we filtered all gene models with hits to InterPro domains that are specific to repetitive elements.

### Phylogenetic reconstruction of Cannabaceae

Multiple sequence alignments were generated using MAFFT (version 7.017)^103^ and phylogenetic analyses were performed using MrBayes (version 3.2.2)^104^. The first phylogenetic reconstruction of the Cannabaceae was based on four markers comprising data from Yang et al. 2013^29^ supplemented with new data generated with primers and protocols published in this manuscript (Supplementary Table 12). Analysis was based on five optimal partitions and models of sequence evolution as estimated by PartitionFinder (version 2.0.0)^105^: atpB-rbcL combined with trnL-F (GTR+I+G); first codon position of rbcL (GTR+I+G); second position of rbcL (SYM+I+G); third position of rbcL (GTR+G); rps16 (GTR+G). An additional phylogenetic reconstruction of the Cannabaceae was based on whole chloroplast genomes (Supplementary Table 12). Analysis was based on eight optimal partitions and models of sequence evolution as estimated by PartitionFinder: tRNA sequence (HKY+I), rRNA sequence (GTR+I), long single copy region (LSC) coding sequence (GTR+I+G), LSC non-coding sequence (GTR+G), short single copy region (SSC) coding sequence (GTR+G), SSC non-coding sequence (GTR+G), inverted repeat region (IR) coding sequence (GTR+G), and IR non-coding sequence (GTR+G). For both Cannabaceae reconstructions additional bootstrap support values were calculated using RAxML (version 8.2.9)^105,106^ using the same partitions applying the GTR+G model. All gene tree reconstructions were based on unpartitioned analysis of protein sequence with the POISSON+G model.

### Orthogroup inference

To determine the relationships between *P. andersonii* and *T. orientalis* genes, as well as with other plant species we inferred orthogroups with OrthoFinder (version 0.4.0)^107^. Since orthogroups are defined as the set of genes that are descended from a single gene in the last common ancestor of all the species being considered, they can comprise orthologous as well as paralogous genes. Our analysis included proteomes of selected species from the Eurosid clade: *Arabidopsis thaliana* TAIR10 (Brassicaceae, Brassicales)^108^ and *Eucalyptus grandis* v2.0 (Myrtaceae, Myrtales) from the Malvid clade^109^; *Populus trichocarpa* v3.0 (Salicaeae, Malpighiales)^110^, legumes *Medicago truncatula* Mt4.0v1^111^ and *Glycine max* Wm82.a2.v1 (Fabaceae, Fabales)^112^, *Fragaria vesca* v1.1 (Rosaceae, Rosales)^48^, *P. andersonii* and *T. orientalis* (Cannabaceae, Rosales) from the Fabid clade (Supplementary Table 7). Sequences were retrieved from phytozome (www.phytozome.net).

### Gene copy number variant detection

To assess orthologous and paralogous relationships between *Parasponia* and *Trema* genes, we inferred phylogenetic gene trees for each orthogroup comprising *Parasponia* and/or *Trema* genes using the neighbour-joining algorithm^113^. Based on these gene trees, for each *Parasponia* gene, its relationship to other *Parasponia* and *Trema* genes was defined as follows. 1) orthologous pair: the sister lineage is a single gene from the *Trema* genome suggesting that they are the result of a speciation event; 2) inparalog: the sister lineage is a gene from the *Parasponia* genome, suggesting that they are the result of a gene duplication event; 3) singleton: the sister lineage is a gene from a species other than *Trema*, suggesting that the *Trema* gene was lost; 4) multi-ortholog: the sister lineage comprises multiple genes from the *Trema* genome, suggesting that the latter are inparalogs. For each *Trema* gene, relationship was defined in the same way but with respect to the *Parasponia* genome (Supplementary Table 8). Because phylogenetic analysis relies on homology we assessed the level of conservation in the multiple-sequence alignments by calculating the trident score using MstatX (https://github.com/gcollet/MstatX)^114^. Orthogroups with a score below 0.1 were excluded from the analysis. Examination of orthogroups comprising >20 inparalogs revealed that some represented repetitive elements; these were also excluded. Finally, orthologous pairs were validated based on the whole-genome alignments used in the annotation reconciliation.

### Assembly of Parasponia and Trema draft genomes

To assess whether gene copy number variants of interest are also present in other, nonreference *Parasponia* and *Trema* genomes, we assembled genomic sequences of *P. rigida*, *P. rugosa*, *T. levigata*, and *T. orientalis* accession RG16 based on the medium-coverage sequence data that was also used for *k*-mer analysis (Supplementary Table 4-5). Assembly was performed with the iterative de Bruijn graph assembler IDBA-UD (version 1.1.1)^115^, iterating from 30-mers (assembling low-coverage regions) to 120-mers (accurately assembling regions of high coverage), with incremental steps of 20. Genes of interest were manually annotated and putatively lost genes or gene fragments were confirmed based on (I.) mapping the medium-coverage reads to the respective *P. andersonii* or *T. orientalis* RG33 reference genome and (II.) genomic alignments (Supplementary Fig. 18–19).

### Nodule-enhanced genes

To assess gene expression in *Parasponia* nodules, RNA was sequenced from the three nodule stages described above as well as uninoculated roots (Supplementary Table 11). RNA-seq reads were mapped to the *Parasponia* reference genome with HISAT2 (version 2.02)^116^ using an index that includes exon and splice site information in the RNA-seq alignments. Mapped reads were assigned to transcripts with featureCounts (version 1.5.0)^117^. Normalization and differential gene expression were performed with DESeq2. Nodule enhanced genes were selected based on >2.0 fold-change and p<=0.05 in any nodule stage compared with uninoculated root controls. Genes without functional annotation or orthogroup membership were excluded. To assess expression of *Parasponia* genes in the hybrid nodules, RNA was sequenced from nodules and uninoculated roots. Here, RNA-seq reads were mapped to a combined reference comprising two parent genomes from *P. andersonii* and *T. tomentosa*.

## Acknowledgments

This work was supported by NWO-NSFC Joint Research project (846.11.005) to WCY, TB and RG, NWO-VICI (865.13.001) to RG, NWO-VENI (863.15.010) to WK, the European Research Council (ERC-2011-AdG294790) to TB and China Scholarship Councils (201303250067) to FB and (201306040120) to DS. We thank Shelley James and Giles Oldroyd for providing germplasm.

All custom scripts and code are available on https://github.com/holmrenser/parasponia_code. The data reported in this paper are tabulated in the Supplementary Materials and archived at NCBI under BioProject numbers PRJNA272473 and PRJNA272482. All analyzed data can be browsed or downloaded through a WebPortal on www.parasponia.org. [For reviewing purposes, an account is available with username *reviewer* and password *D7yGNEkNv25e*].

## Contributions

This research was led by RG, who together with TB conceived the project. *Trema orientalis* accessions, including rhizosphere samples, were collected in Sabah Parks (Malaysian Borneo) by RG in an expedition organised by MS and RR. FACS studies to estimate genome sizes were done by FB and RHe. Plant propagations and tissue isolations were done by FB, WL, QC, TS, DS, YR, MH, WY and RG. Arbuscular mycorrhiza assays were done by TS, YR and WK. Studies on hybrid plants were done by FB, QC, DJvdH and EF, and ARA assays by FB and EF. Light and electron microscopy studies were conducted by FB and EF. DNA and RNA was isolated by JV, JH and WL, and sequencing was done by ES. Chloroplast analysis was conducted by RvV, RHo and BG, Bioinformatic analyses were done by RvV, RHo, LS, JJ and SS, and manual curations by RvV, RHo, LR, AvZ, TAKW, JJ, KM, WK, and RG. RvV, RHo, LR, MES, TB, SS, and RG wrote the manuscript.

---

**Supplementary Figure 1:**
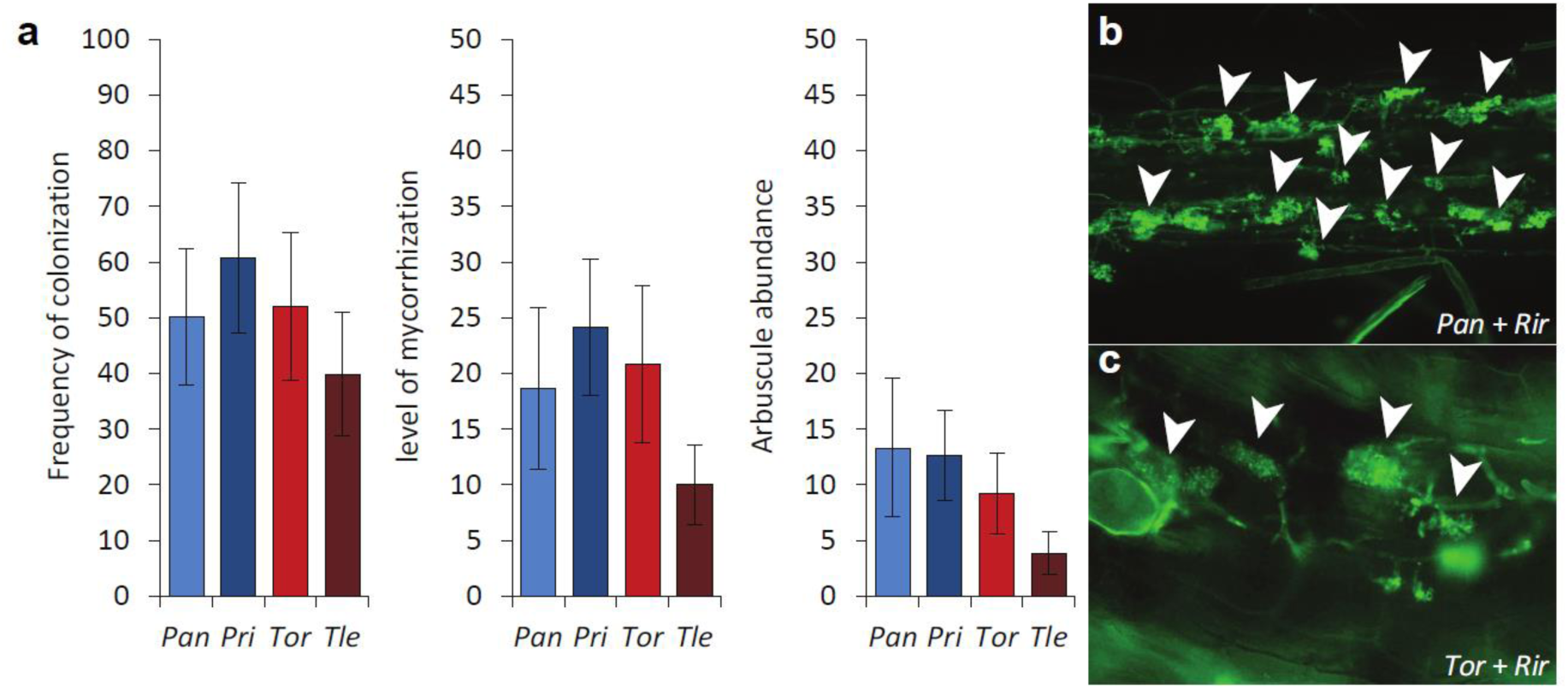
Arbuscular mycorrhization of *Parasponia* and *Trema* species. **(a)** Mycorrhization efficiency of *Parasponia andersonii* WU01.14 (Pan), *Parasponia rigida* WU20 (Pri), *Trema orientalis* RG33 (Tor) and *Trema levigata* WU50 (Tle), 6 weeks post inoculation with *Rhizophagus irregularis* (*Rir*, n=10, error bars denote standard errors). **(b, c)** Confocal image of WGA-Alexafluor 488-stained arbuscules in root segment of either *P. andersonii* (Pan) **(b)** or *T. orientalis* (Tor) **(c)**.

**Supplementary Figure 2:**
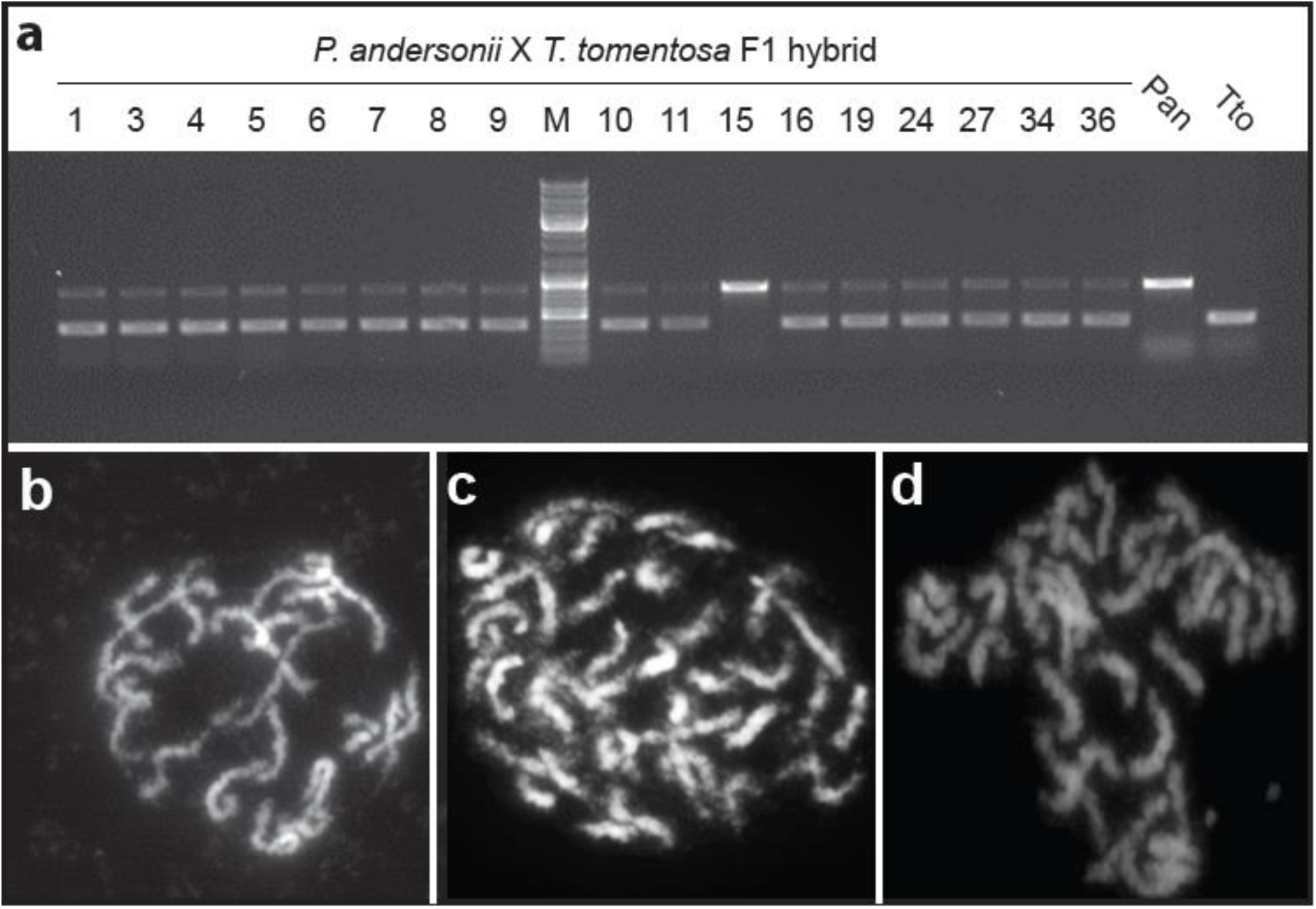
Genotyping of *Parasponia andersonii* × *Trema tomentosa* F_1_ hybrid plants. **(a)** Genotyping of 17 putative F_1_ hybrid plants of the cross *P. andersonii* (Pan) × *T. tomentosa* (Tto) using amplified length polymorphism due to an indel in the *LAX1* promoter. M: generuler DNA ladder mix (Fermentas). Hybrid plants 4, 8, 9, 16, 19 and 36 were used for further experiments. **(b-d)** Mitotic metaphase chromosome complement of *P. andersonii* (2n=2×=20) **(b)**, *T. tomentosa* (2n=4×=40) **(c)**, and *P. andersonii × T. tomentosa* F1 hybrid (2n=3x=30) **(d)**.

**Supplementary Figure 3:**
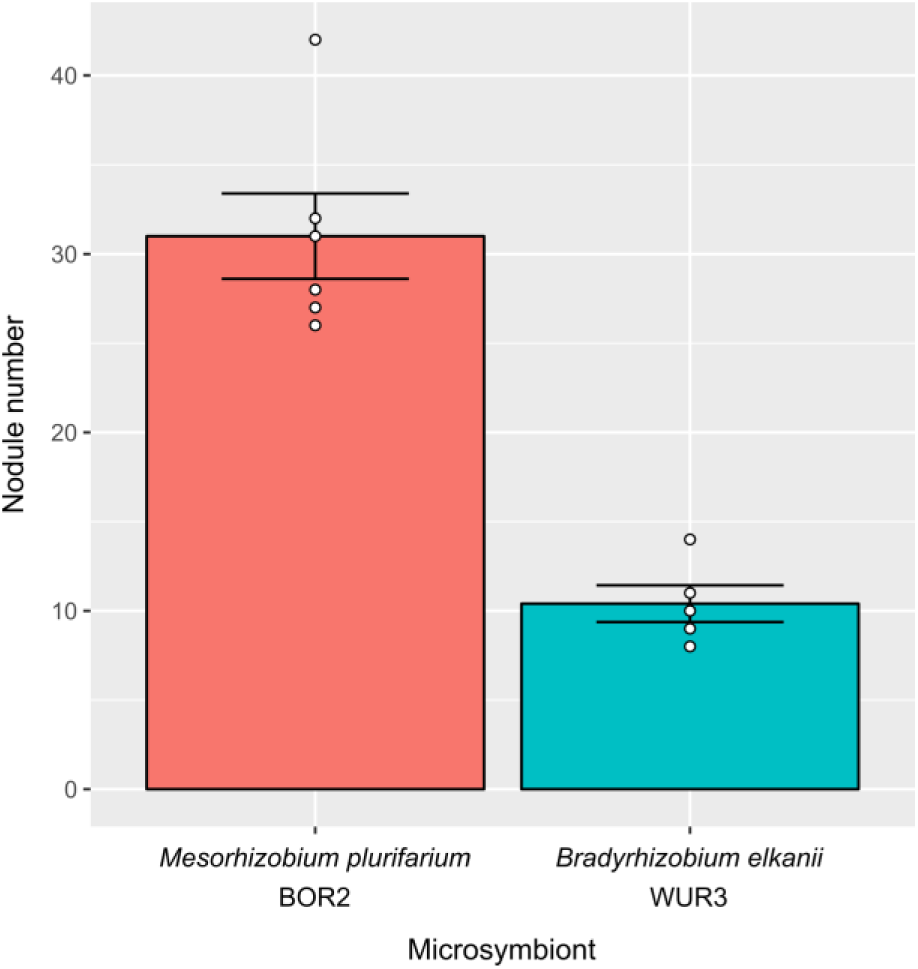
Nodulation efficiency of *Parasponia andersonii*. Mean number of nodules on roots of *P. andersonii* inoculated with either *Mesorhizobium plurifarium* BOR2 (n=6) or *Bradyrhizobium elkanii* WUR3 microsymbionts (n=5) (6 weeks post inoculation). Dots represent individual measurements.

**Supplementary Figure 4:**
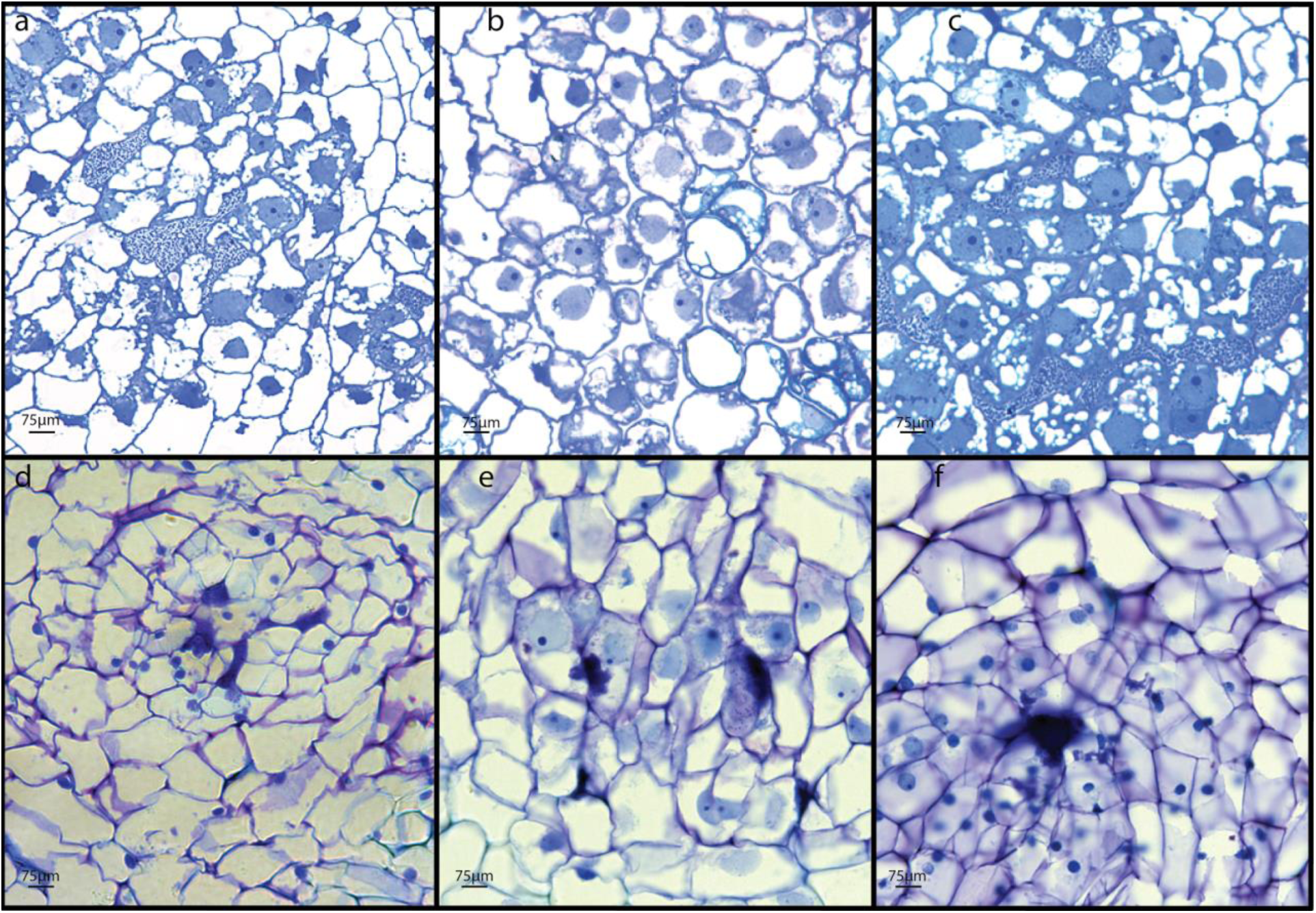
Longitudinal sections of root nodules of *Parasponia andersonii* × *Trema tomentosa* F1 hybrid plants. Hybrid plants H4, H8, H9, H16, H19 and H36 were clonally propagated and inoculated and inoculated with either *Bradyrhizobium elkanii* WUR3 **(a-c)** or *Mesorhizobium plurifarium* BOR2 **(d-f)**. **(a)** H4 nodule induced by *B. elkanii* WUR3. **(b)** H8 nodule induced by *B. elkanii* WUR3. **(c)** H9 nodule induced by *B. elkanii* WUR3. **(d)** H16 nodule induced by *M. plurifarium* BOR2. **(e)** H19 nodule induced by *M. plurifarium* BOR2. **(f)** H36 nodule induced by *M. plurifarium* BOR2. Note absence of intracellular infection in all sectioned nodules.

**Supplementary Figure 5:**
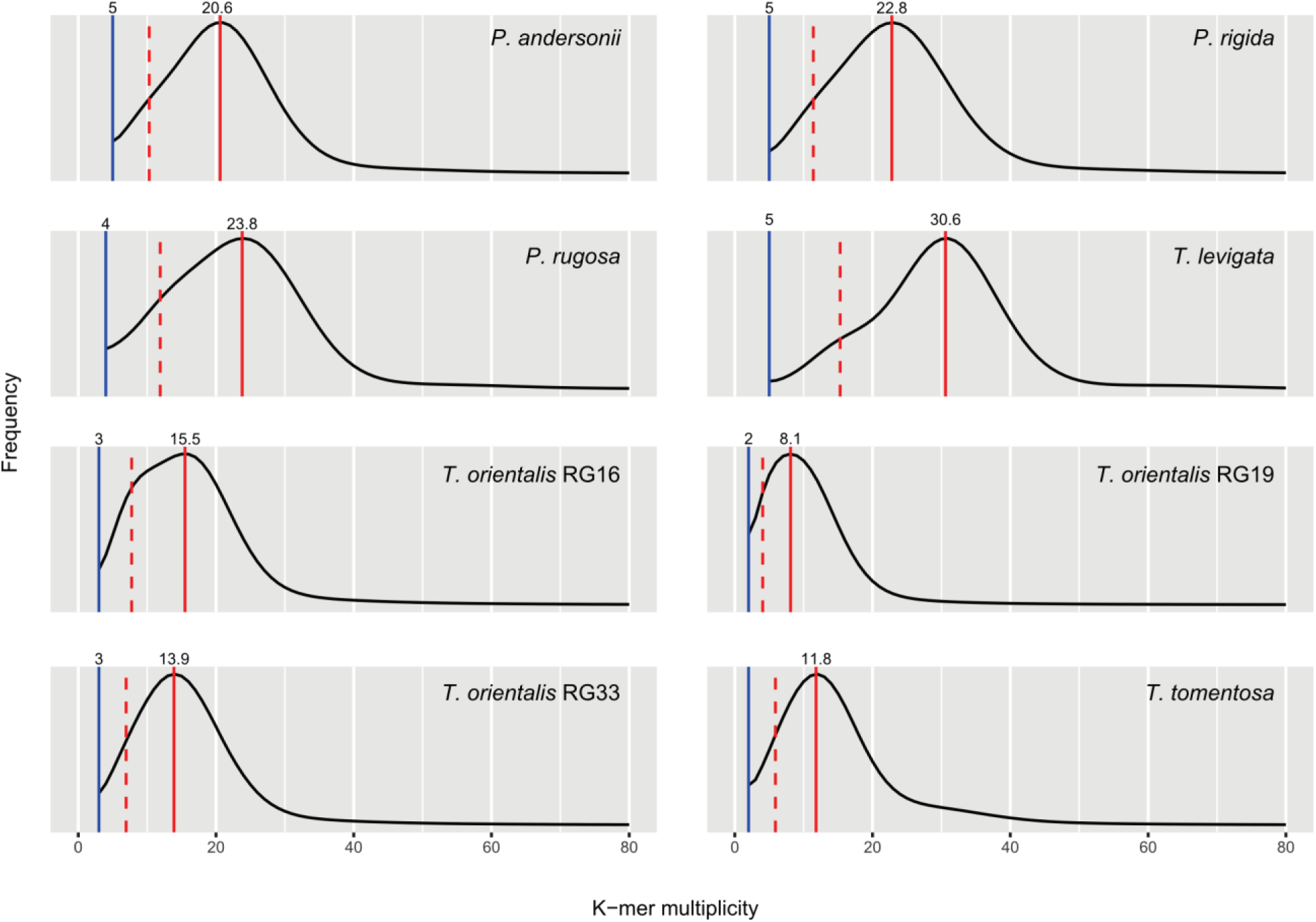
Genome coverage and heterozygosity estimates based on k-mer analysis of *Trema* and *Parasponia* species. Plots of 21-mer multiplicity frequencies based on jellyfish output showing that *T. levigata* and *T. orientalis* RG16 are relatively heterozygous. Solid red lines indicate estimated genome coverage corresponding to homozygous sequence; dashed red lines indicate half the estimated genome coverage corresponding to heterozygous sequence; blue lines indicate estimated error multiplicity threshold.

**Supplementary Figure 6:**
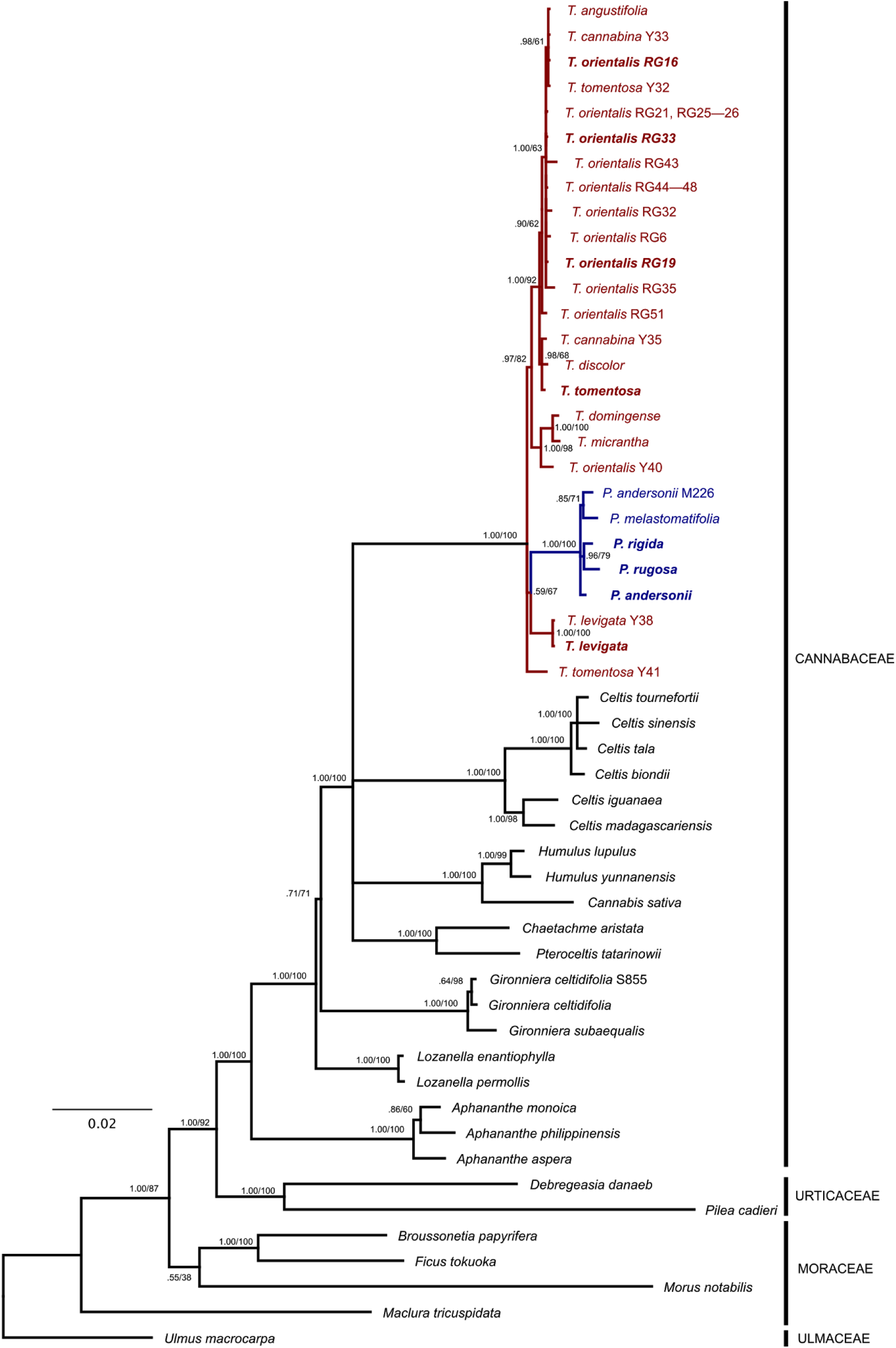
Phylogenetic reconstruction of the Cannabaceae based on combined analysis of four plastid markers. Node values indicate posterior probability / RAxML bootstrap support; scale bar represents substitutions per site. *Parasponia* lineage is in blue, *Trema* lineages are in red. Note that sister relationship of *Parasponia* and *T. levigata* has low bootstrap support, but is independently supported by four shared sequence insertions (Supplementary Fig. 8). Accessions selected for comparative genome analysis in bold. GenBank accession numbers are in Supplementary Table 12.

**Supplementary Figure 7:**
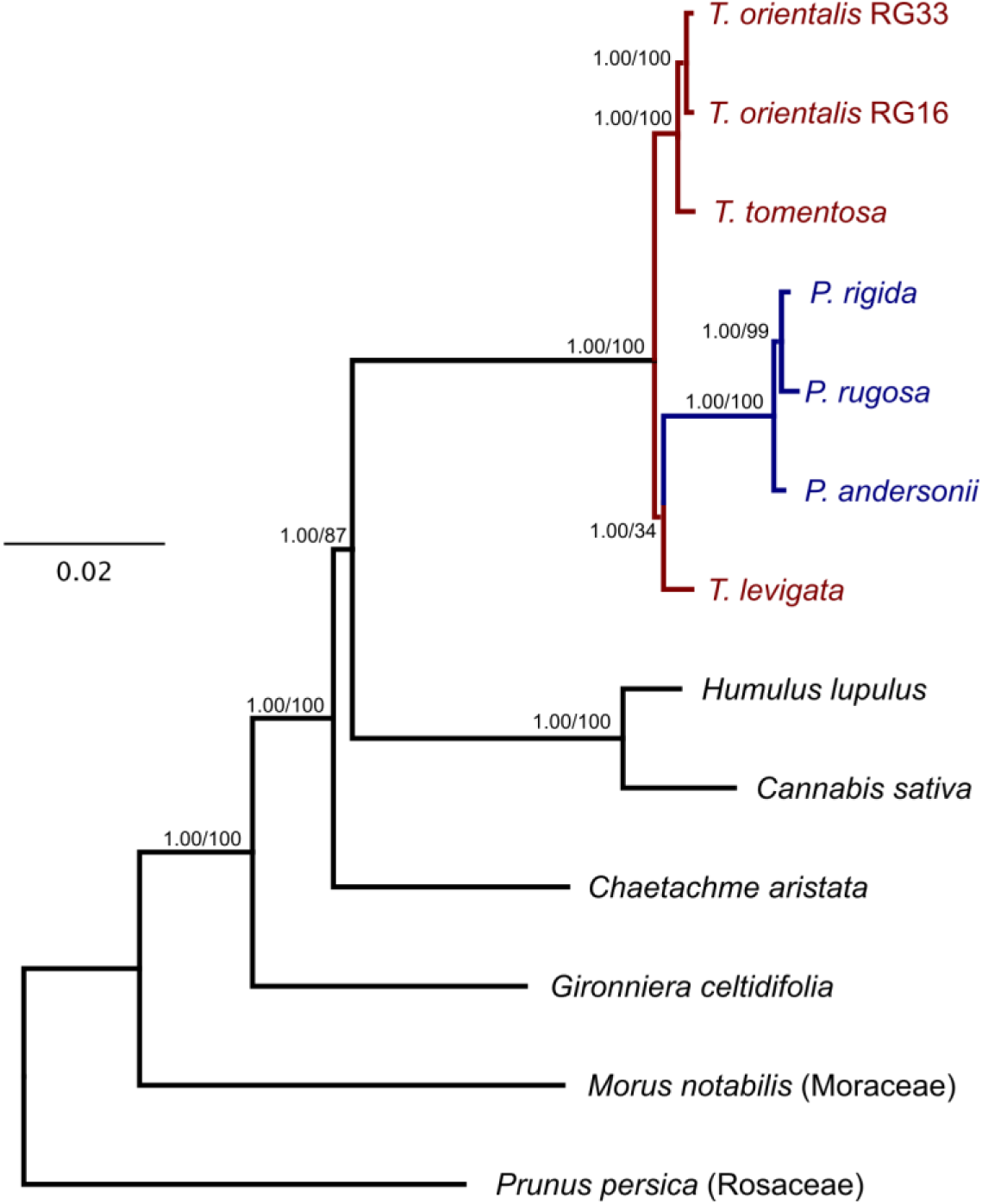
Phylogenetic reconstruction of the Cannabaceae based on chloroplast genomes. Bayesian tree based on a combined analysis of eight data partitions (see Methods). *Parasponia* lineage is in blue, *Trema* lineages are in red. Note that sister relationship of *Parasponia* and *T. levigata* has low bootstrap support but is independently supported by fou rshared sequence insertions (Supplementary Fig. 8). Node values indicate posterior probability / RAxML bootstrap support; scale bar represents substitutions per site. GenBank accession numbers are in Supplementary Table 12.

**Supplementary Figure 8:**
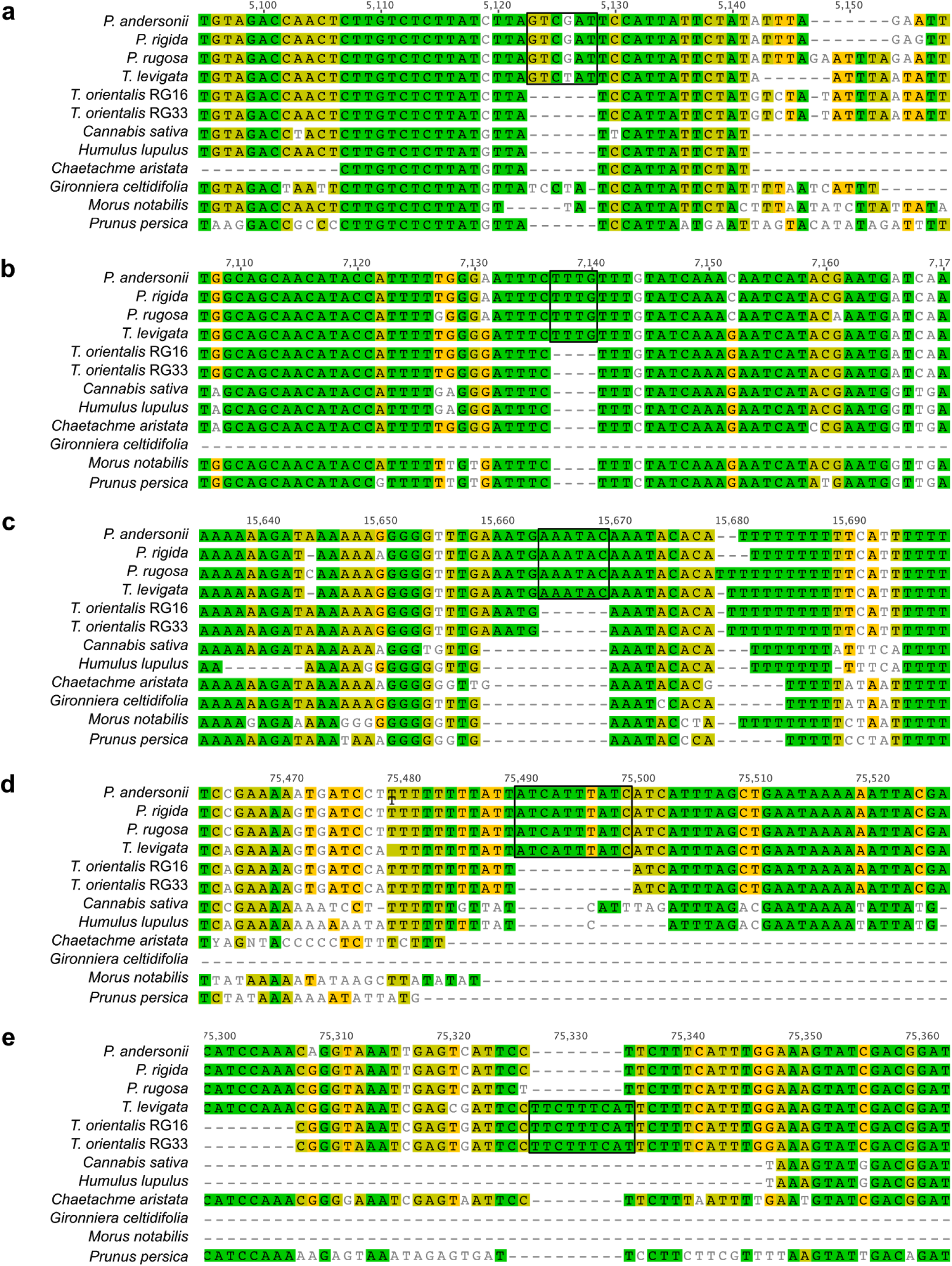
Chloroplast genome insertions in Cannabaceae. Shared sequence insertions in chloroplast genomes supporting **(a-d)** or refuting **(e)** sister relationship of *Parasponia* and *Trema levigata*. **(a)** *matK*-*rps16* intergenic spacer, **(b)** *rps16*-*psbK* intergenic spacer, **(c)** *atpF* intron, **(d, e)** *petA*-*psbJ* intergenic spacer. Numbers indicate alignment coordinates; colours indicate percent identity while ignoring gaps: green = 100%, olive = 80-100%, yellow = 60-80%; black rectangles mark shared sequence insertions concerned.

**Supplementary Figure 9:**
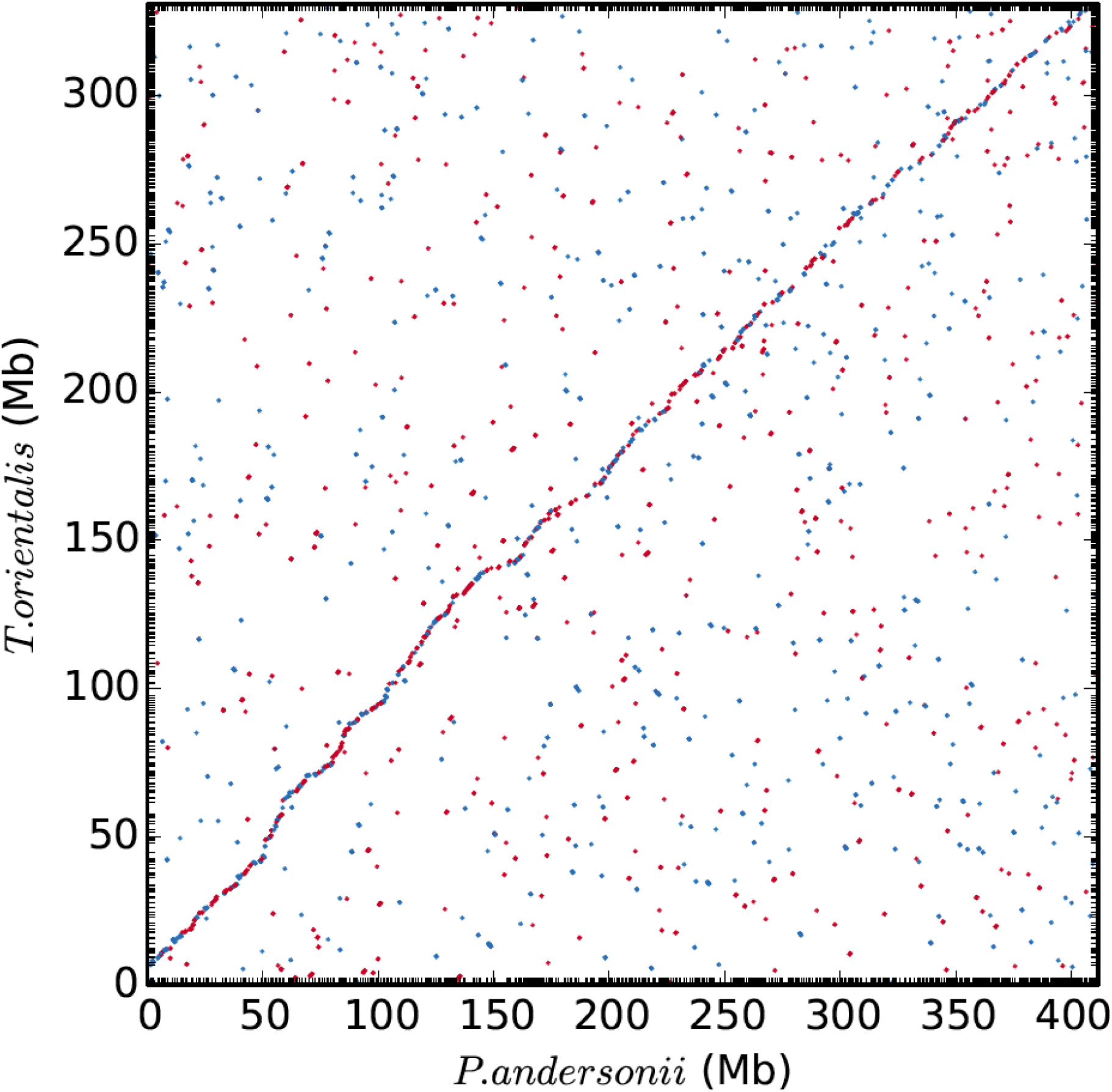
Whole genome alignment dotplot for *P. andersonii* and *T. orientalis* RG33. Maximal unique matching (MUM) alignments were generated using nucmer 4.0.0beta with the following settings: breaklength 500, mincluster 200, maxgap 100, minmatch 80, minalign 7000. Forward alignments are red, reverse alignments are blue. Scaffolds are ordered by alignment size, which results in a clear diagonal line indicating the collinearity of the two genomes.

**Supplementary Figure 10:**
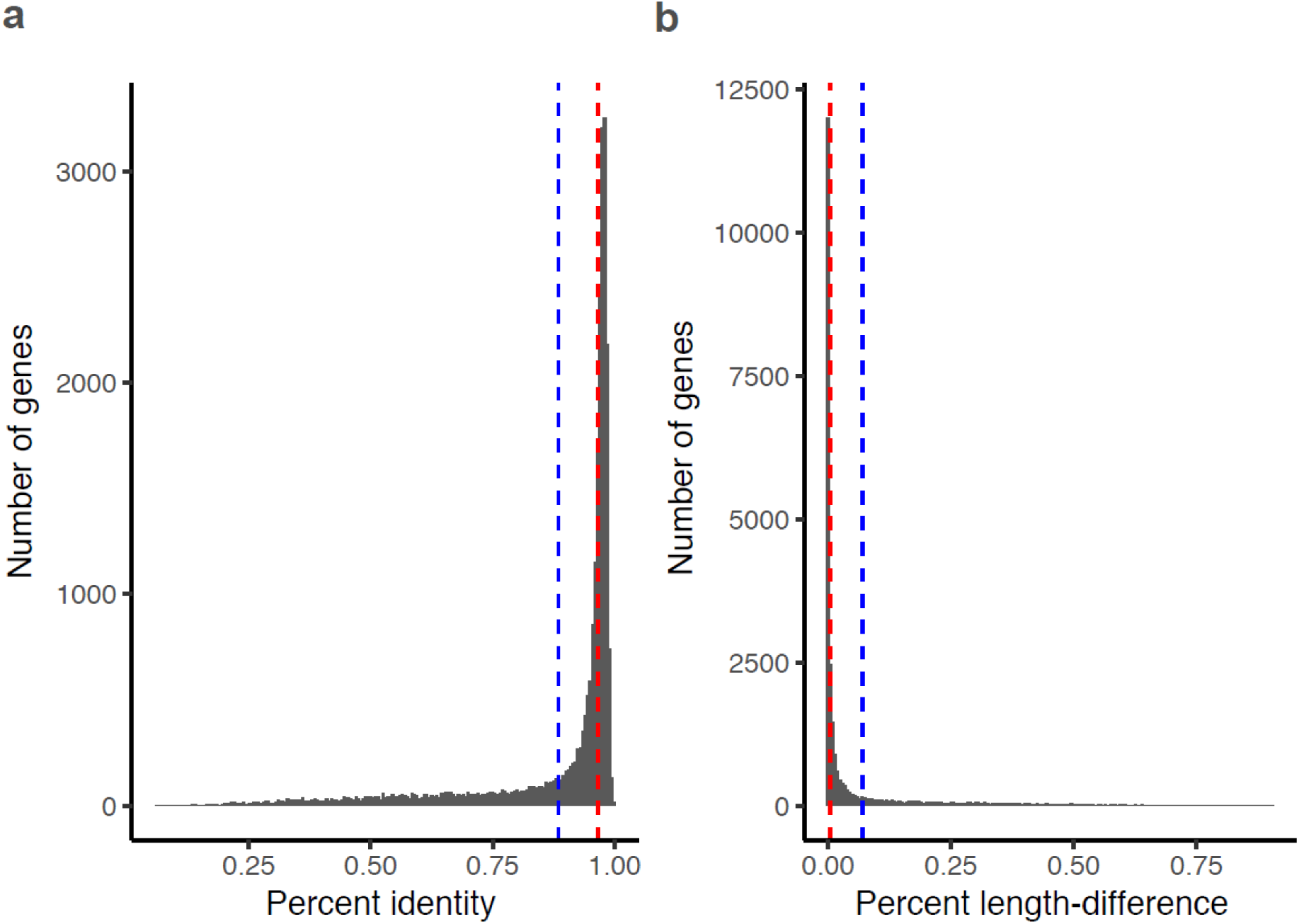
Identity of *P. andersonii* - *T. orientalis* putative orthologous gene pairs. Histograms of **(a)** percent nucleotide identity (calculated by taking the fraction of identical nucleotides ignoring end gaps using global alignments produced by MAFFT version 7.017^103^) and **(b)** length difference of all 25,605 orthologous gene pairs from *P. andersonii* and *T. orientalis* as a percentage of the longest gene. Red line indicates median, blue line indicates mean.

**Supplementary Figure 11:**
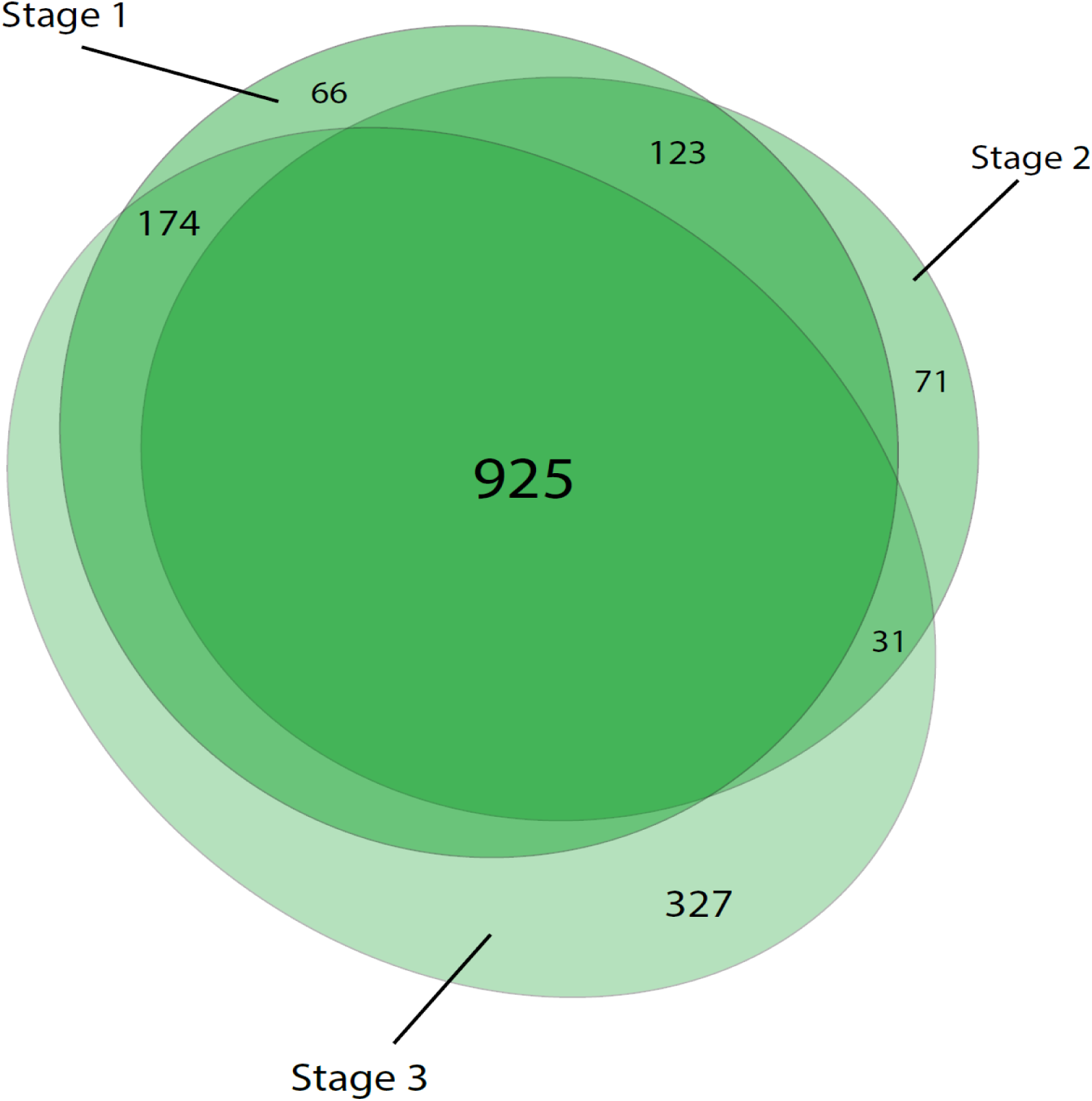
Venn diagram of *P. andersonii* nodule enhanced genes in 3 developmental stages. Nodule developmental stages according to Fig. 1h-j. List of genes is given in Supplementary Table 9. *Parasponia andersonii* genes are considered ‘nodule enhanced’ when expression is increased >2-fold in any of 3 nodule developmental stages when compared to non-inoculated root sample. Largest fraction concerns genes enhanced in all 3 stages.

**Supplementary Figure 12:**
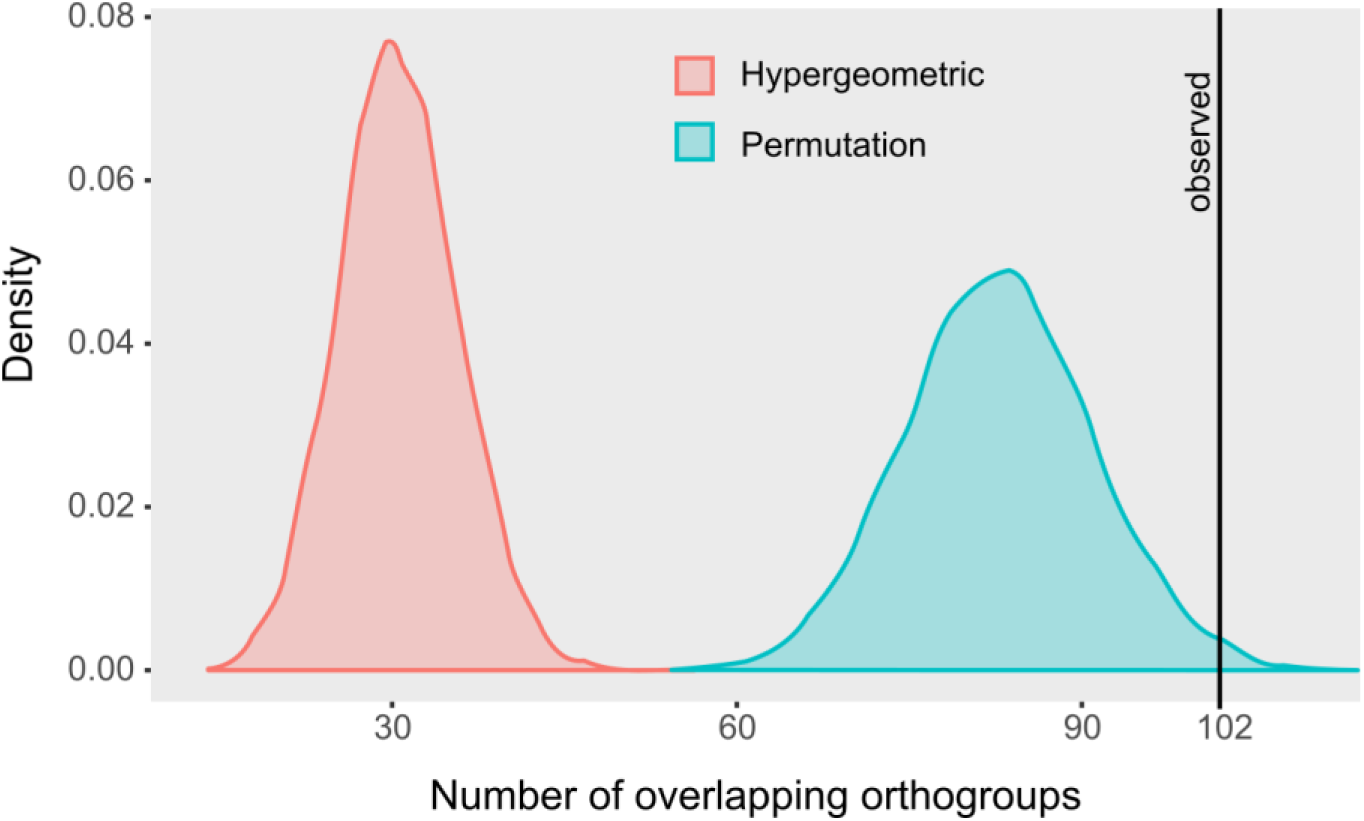
Statistical testing of common utilization of genes in *Parasponia* and medicago. To assess common utilization of genes in *Parasponia* and medicago nodules we performed statistical testing of overlap between *Parasponia andersonii* and medicago nodule-enhanced genes. Overlap was calculated based on orthogroup membership (i.e. when an orthogroup contains nodule-enhanced genes from *P. andersonii* and medicago it is scored as overlap). Significance of set overlaps is usually calculated based on the hypergeometric distribution. However, because larger orthogroups have higher chance of overlap, the hypergeometric is not suitable. We therefore assessed significance with a permutation test where the null distribution is based on overlap found when gene-orthogroup membership is randomized (n=10,000).Figure shows density plots of both hypergeometric distribution and permutation random variates. Vertical line shows the observed number of 102 overlapping orthogroups (p<0.02 based on permutation test).

**Supplementary Figure 13:**
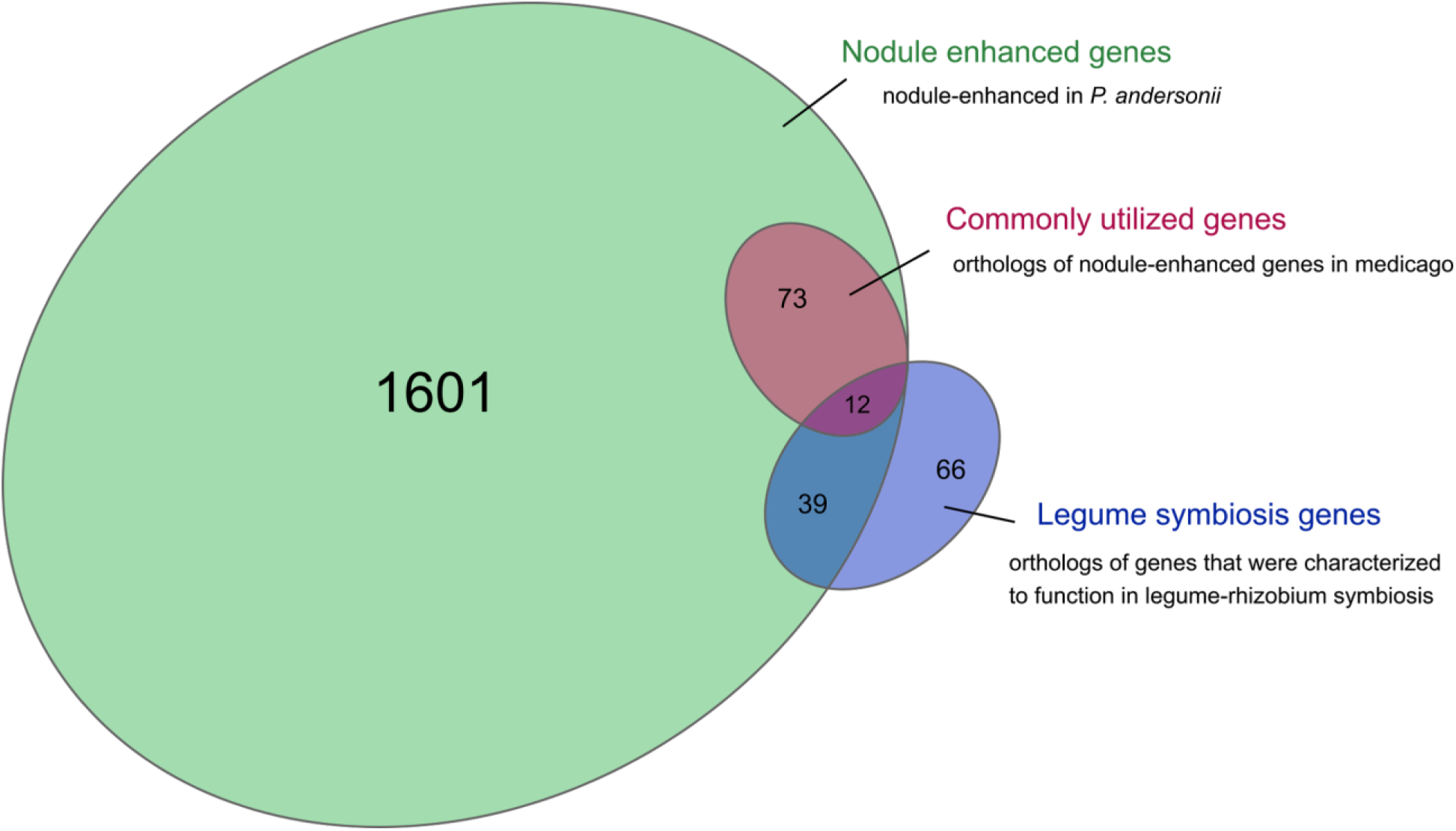
Venn diagram of *P. andersonii* symbiosis gene sets. Nodule enhanced genes have a significantly enhanced expression level (fold change > 2, p < 0.05, DESeq2 Wald test) in any of three developmental stages (N = 1725; Supplementary Fig. 11; Supplementary Table 9). Commonly utilized genes are nodule-enhanced in *P. andersonii* as well as in the legume medicago^31^ (N = 85; Supplementary Table 10, Supplementary Data File 2). Legume symbiosis genes are orthologs of genes that were characterized to function in legume-rhizobium symbiosis (N = 117; Supplementary Table 1, Supplementary Data File 1).

**Supplementary Figure 14:**
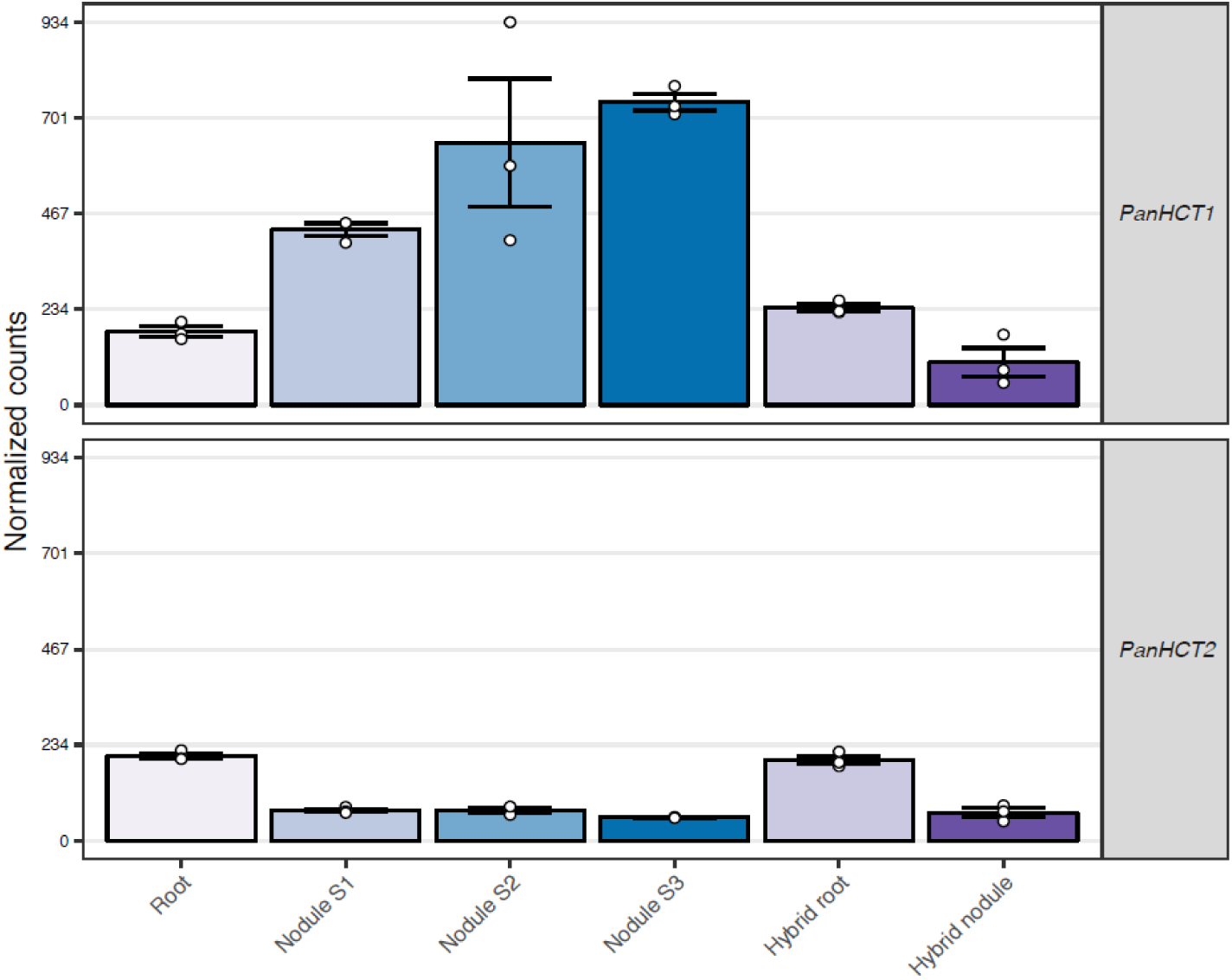
Expression profile of *PanHCT1* and *PanHCT2* genes. Expression of *P. andersonii HYDROXYCINNAMOYL*-*COA SHIKIMATE TRANSFERASE 1* (*PanHCT1*) and *PanHCT2* in *P. andersonii* roots, stage 1-3 nodules, and in *P. andersonii* × *T. tomentosa* F_1_ hybrid roots and nodules (line H9). *PanHCT1* and *PanHCT2* represent the only *Parasponia*-specific gene duplication in the defined symbiosis gene set, as *PanHCT1* is upregulated in nodules. Expression is given in DESeq2 normalized read counts, error bars represent standard error of three biological replicates, dots represent individual expression levels.

**Supplementary Figure 15:**
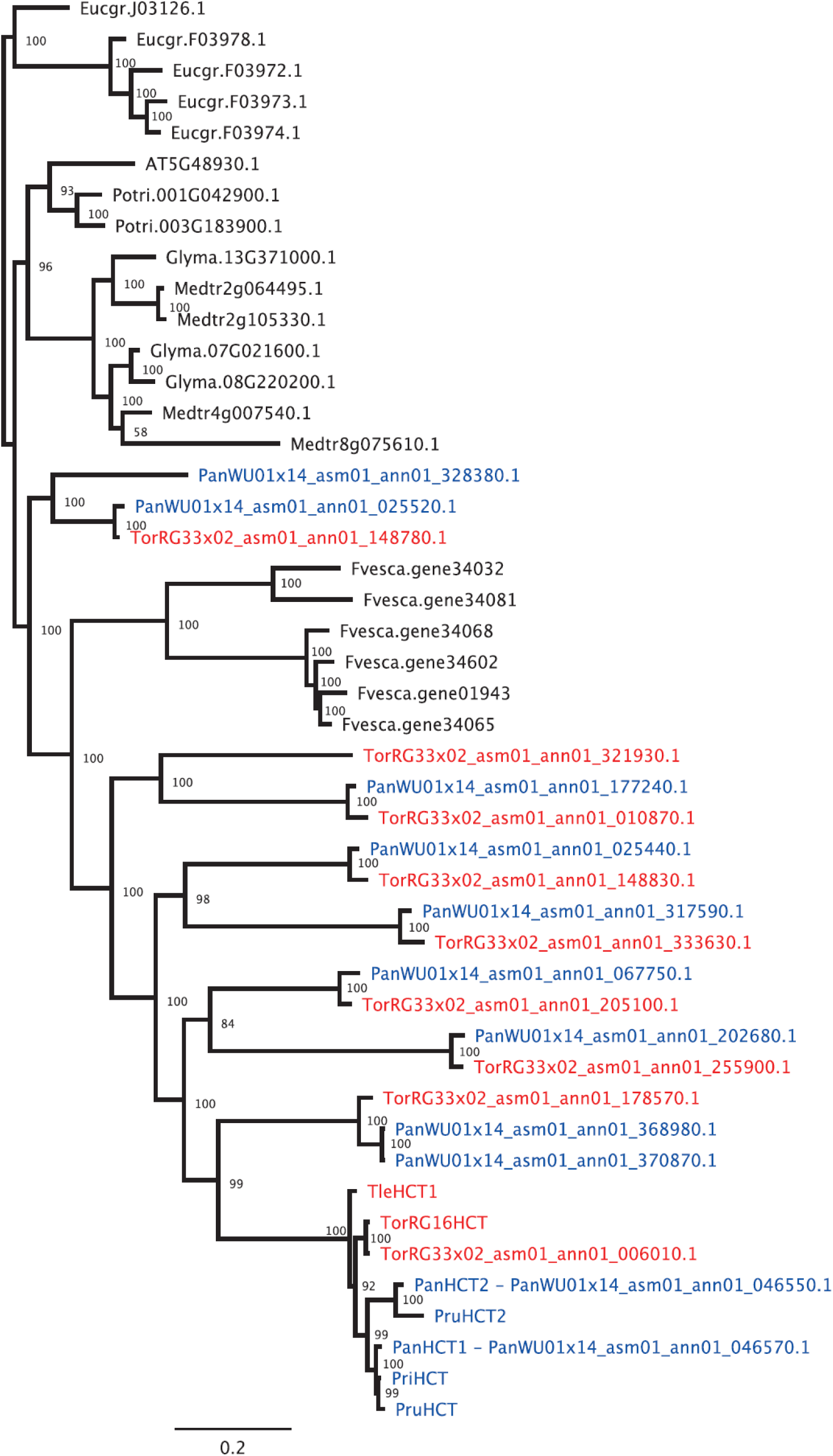
Phylogenetic reconstruction of Hydroxycinnamoyl-CoA Shikimate Transferase (HCT) orthogroup. *HCT* orthogroup was created by merging OG0001291, OG0016758, OG0016791, OG0018560, OG0020327, OG0020921,
OG0022256 & OG0023772, supplemented with HCT1 and HCT2 orthologs of *P. rigida*, *P. rugosa*, *T. orientalis* RG16 and *T. levigata. PriHCT2* is a putative pseudogene and was not included. HCT1 and HCT2 represent the only *Parasponia* specific gene duplication in the defined symbiosis gene set, as *PanHCT1* was found to be upregulated in nodules. Species included: *Parasponia andersonii* (Pan); *P. rigida* (Pri); *P. rugosa* (Pru) (all in blue); *Trema orientalis* (Tor); *T. orientalis* RG16 (TorRG16); *T. levigata* (Tle) (all in red); *Medicago truncatula* (Mt); *Glycine max* (Glyma), *Populus trichocarpa* (Potri); *Fragaria vesca* (Fvesca); *Eucalyptus grandis* (Eugr); *Arabidopsis thaliana* (AT). Phylogenetic inference was calculated using MrBayes 3.2.2. Scale bar represents substitutions per site.

**Supplementary Figure 16:**
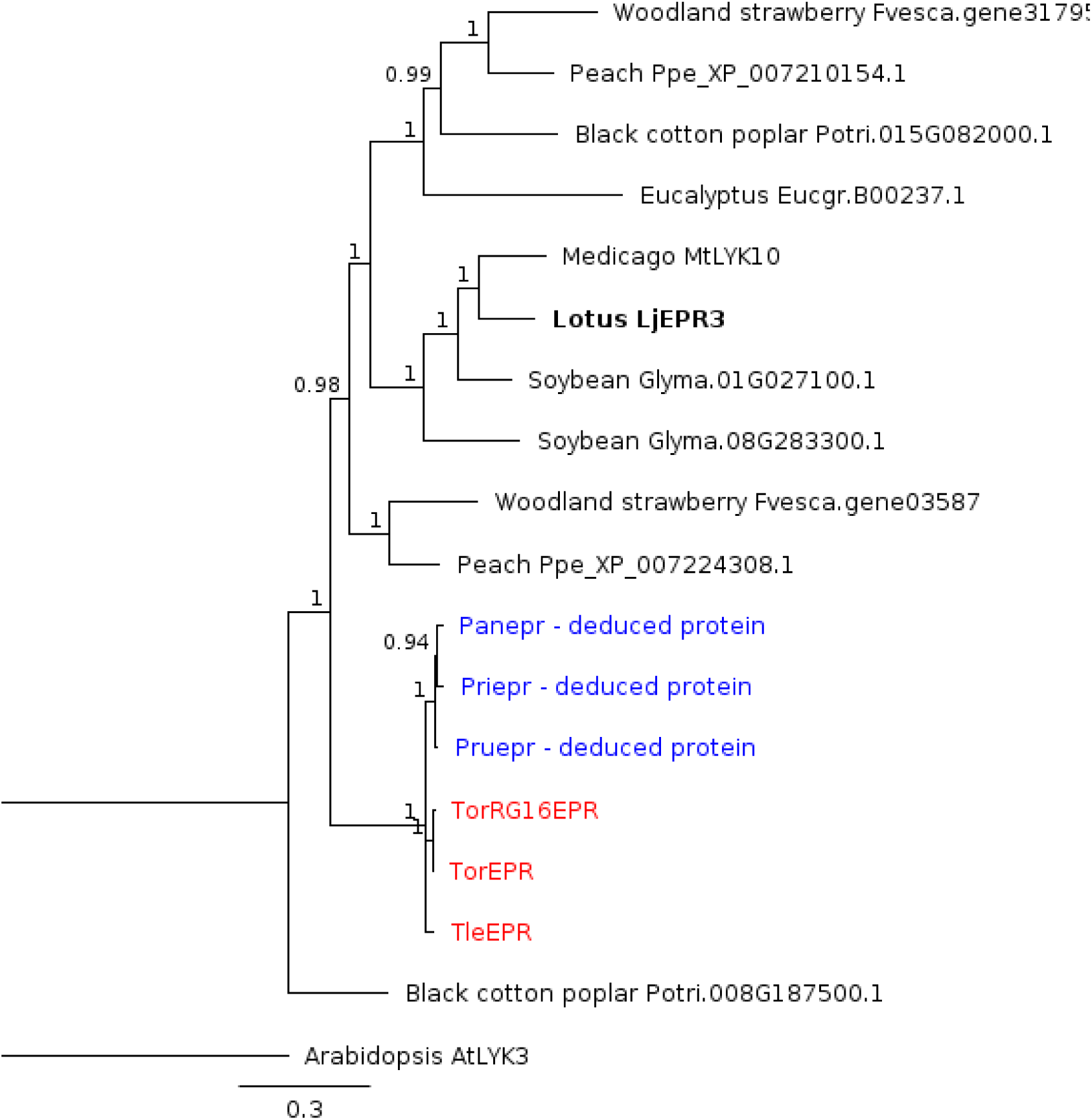
Phylogenetic reconstruction of the EPR3 orthogroup. Alignment of orthogroup OG0010070 containing exopolysaccharide receptor LjEPR3. Note that all *Parasponia* species lack a functional *EPR* (Supplementary Fig. 15). Species included: *Trema orientalis* RG33 (Tor); *Trema orientalis* RG16 (TorRG16); *Trema levigata* (Tle) (all in red); *Parasponia Andersonii* (Pan); *Parasponia Rigida* (Pri) *Parasponia Rugosa* (Pru) (all in blue). *Medicago truncatula* (Mt); *Glycine max* (Glyma), *Populus trichocarpa* (Potri); *Fragaria vesca* (Fvesca); *Eucalyptus grandis* (Eugr). Phylogenetic inference was calculated using MrBayes 3.2.2. Scale bar represents substitutions per site.

**Supplementary Figure 17:**
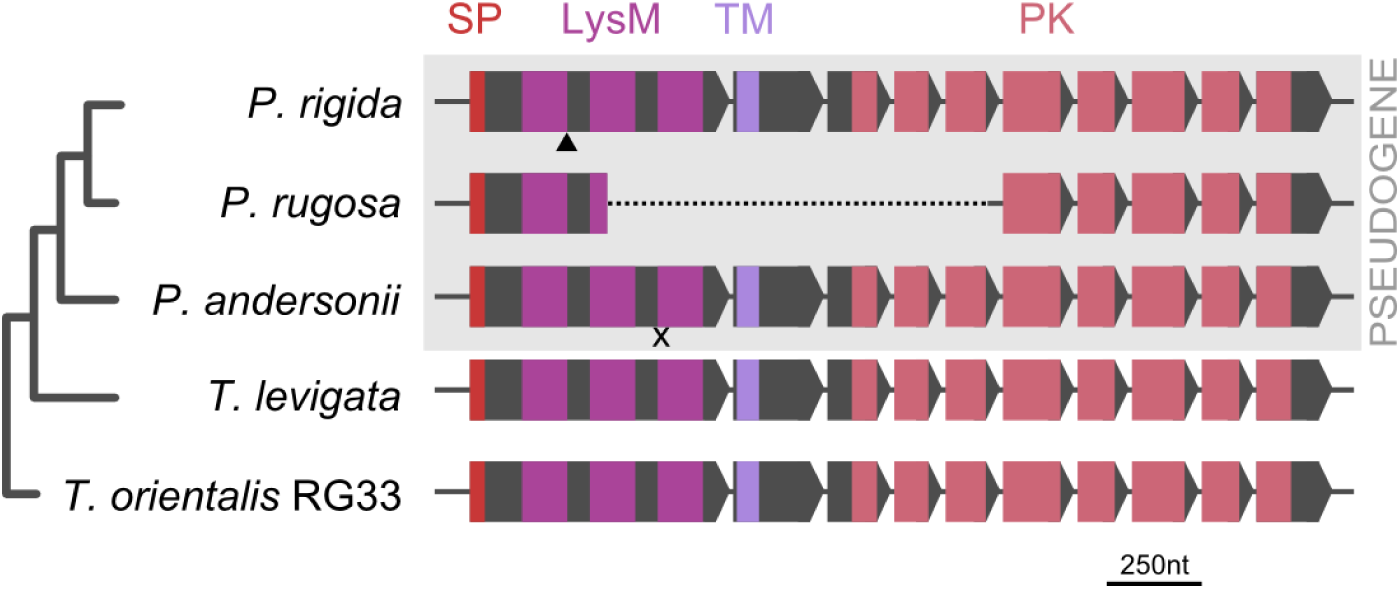
Independent pseudogenization in *Parasponia* species of *EPR* that is orthologous to the *Lotus japonicus* exopolysaccharide receptor *LjEPR3*. Introns are indicated, but not scaled. X indicates premature stop codon in *P. andersonii epr*, triangle indicate frame-shift in *P. rigida epr*, whereas *P. rugosa epr* contains a large deletion. SP = signal peptide (red); LysM: 3 Lysin Motif domains (magenta); TM = transmembrane domain (lilac); PK = protein kinase (pink).

**Supplementary Figure 18:**
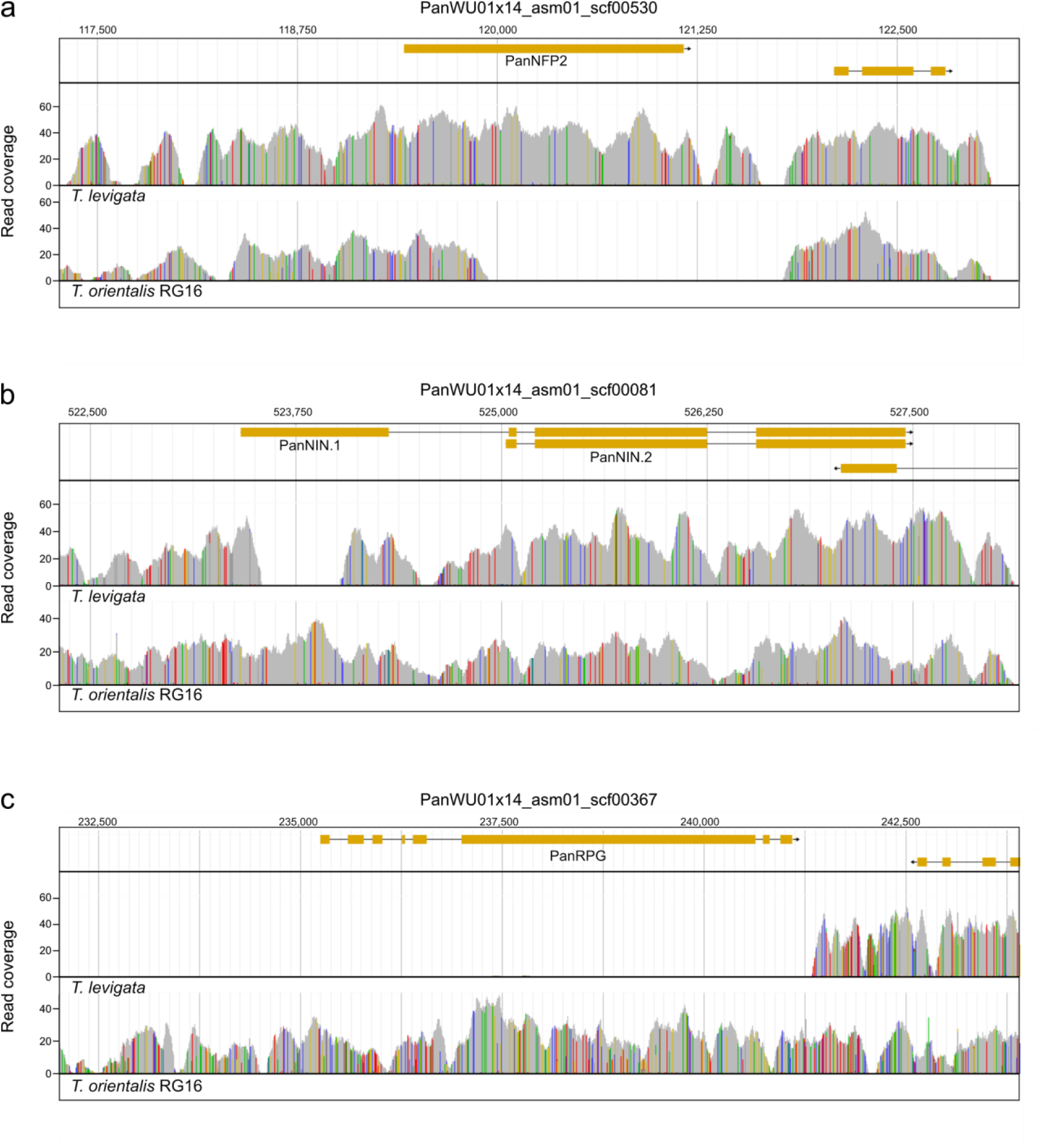
Read mappings of *Trema orientalis* RG16 and *T. levigata* to the *Parasponia andersonii* genome. Read mappings to gene region of **(a)** *PanNFP2*, illustrating absence of a large part of the gene in *T. orientalis* RG16, **(b)** *PanNIN*, illustrating absence of a large part of the canonical first exon in *T. levigata*, **(c)** *PanRPG*, illustrating absence of the gene in *T. levigata*. Coordinates on the x-axis correspond to those of the *P. andersonii* scaffold; orange bars depict *P. andersonii* gene models; histograms depict read coverage in grey; nucleotide differences from the *P. andersonii* reference scaffold are in color (green = adenine, blue = cytosine, yellow = guanine, red = thymine).

**Supplementary Figure 19:**
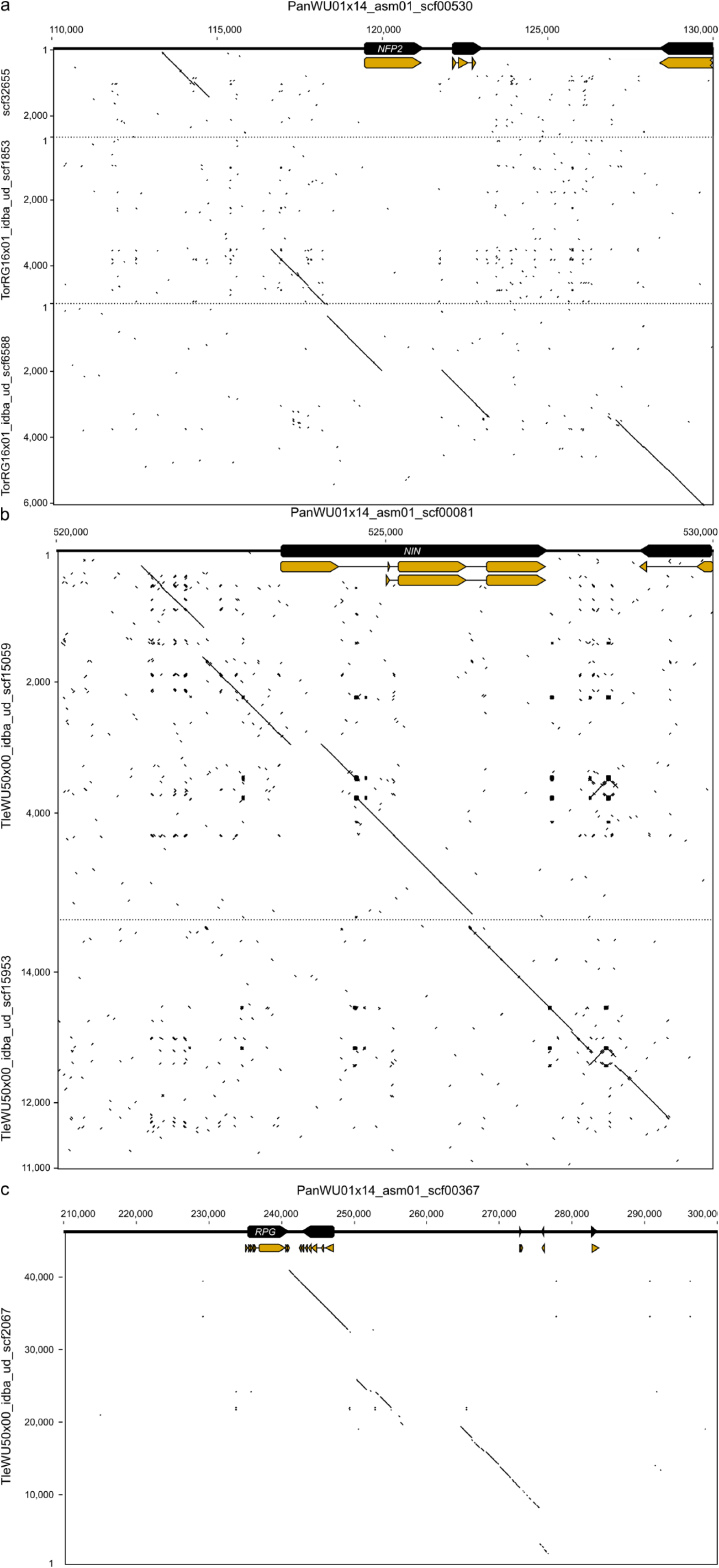
Genomic alignments of *Trema orientalis* RG16 or *Trema levigata* to *Parasponia andersonii NFP2*, *NIN*, *and RPG* gene regions. Genome alignment(s) of **(a)** *T. orientalis* RG16 with *PanNFP2* gene region, **(b)** *T. levigata* with *PanNIN* gene region, **(c)** *T. levigata* with *PanRPG* gene region. Coordinates correspond to those on the draft genome scaffolds; *Parasponia andersonii* gene and CDS models are depicted in black and orange, respectively; different genomic scaffolds are separated by dashed lines.

**Supplementary Figure 20:**
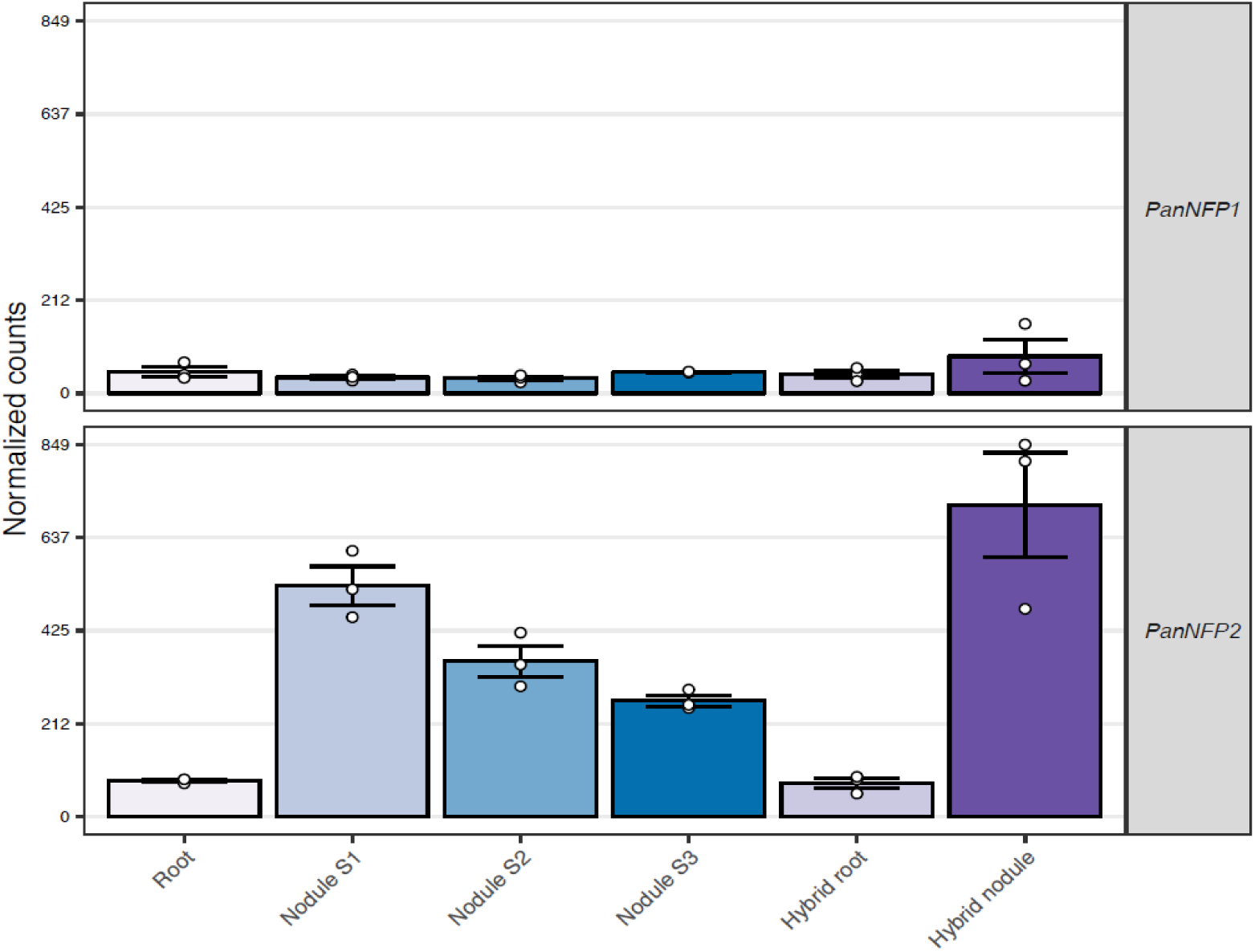
Expression profile of *PanNFP1* and *PanNFP2* genes. Expression of *P. andersonii NOD FACTOR PERCEPTION 1* (*PanNFP1*) and *PanNFP2* in *P. andersonii* roots, stage 1-3 nodules, and in *P. andersonii × T. tomentosa* F1 hybrid roots and nodules. Expression is given in DESeq2 normalized read counts, error bars represent standard error of three biological replicates, dots represent individual expression levels.

**Supplementary Figure 21:**
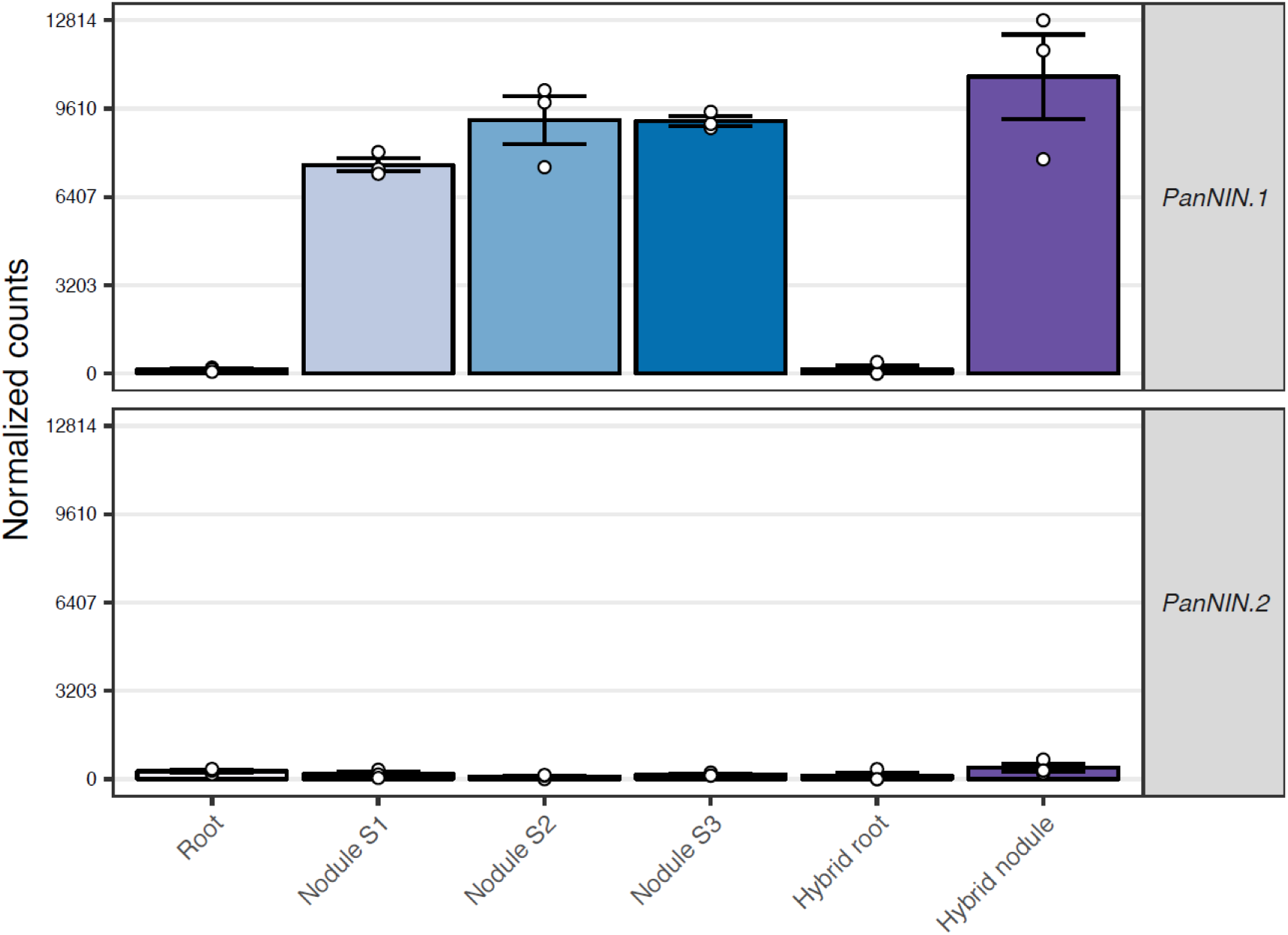
Expression of *P. andersonii NODULE INCEPTION (PanNIN)* gene splice variants. *PanNIN.1* encodes a canonical symbiotic protein, whereas *PanNIN.2* encodes a shorter protein variant that is the result of an alternative start site in an intron. Expression levels were determined by identifying unique DNA sequences for both variants; spanning the intron in case of *PanNIN.1* (CTGCCAAGCGCTTGAGGCTGTTGATCTT), or including the start site of *PanNIN.2* (GCCAATTACCTTGCAGGCTGTTGATCTT) and counting all occurrences in the RNA-seq reads. DESeq2 size factors were used to normalize these counts. The fraction of these normalized counts between *PanNIN. 1* and *PanNIN.2* was used to scale the expression levels. Error bars represent standard error of three biological replicates, dots represent individual expression levels.

**Supplementary Figure 22:**
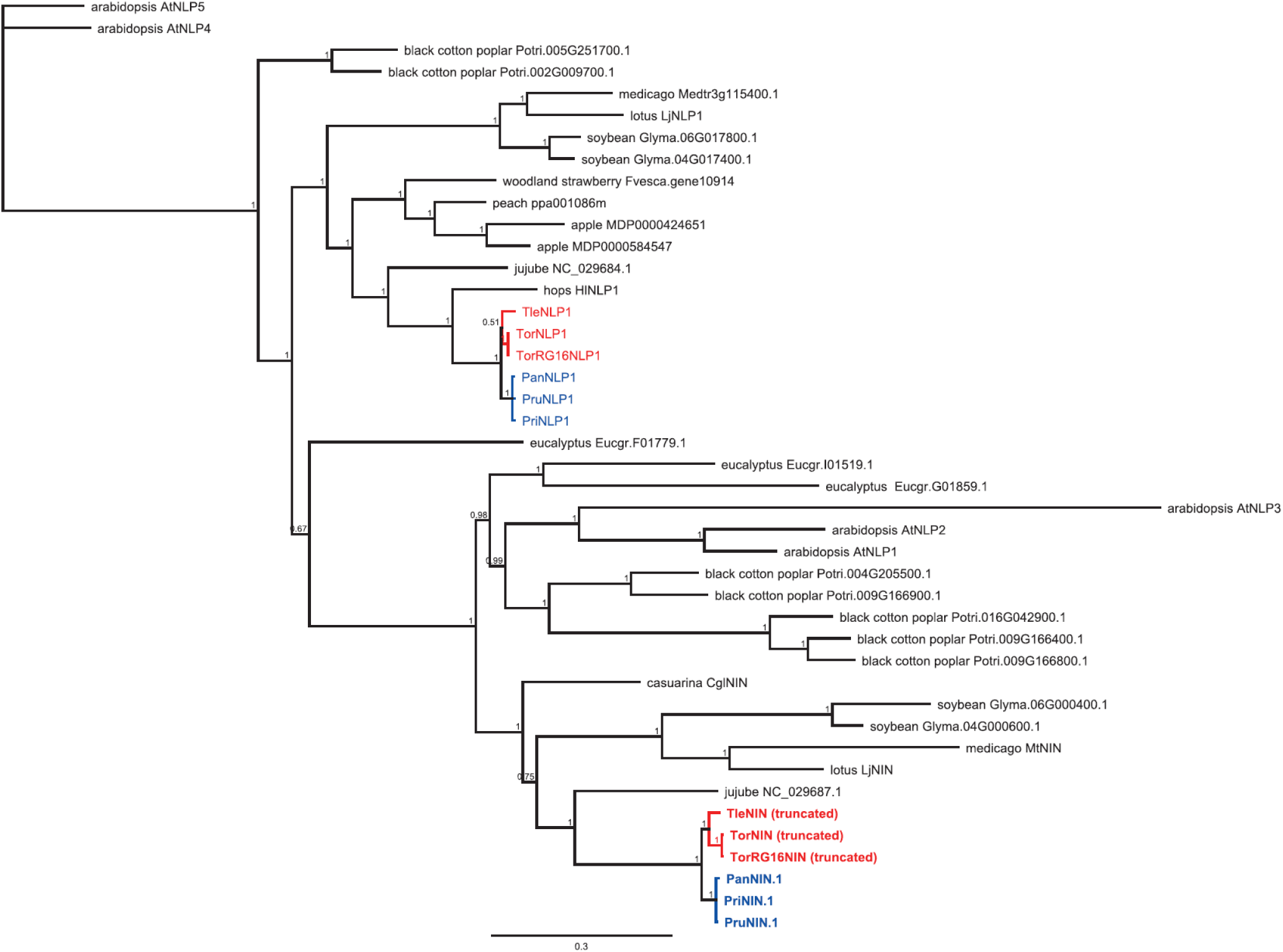
Phylogenetic reconstruction of NIN orthogroup. Alignment of OG0001118, which includes NIN and NLP1 (NIN-LIKE PROTEIN 1)-like proteins, supplemented with additional species. AtNLP4 and AtNLP5 were included as outgroup. *Parasponia* spp. marked in blue, *Trema* spp. In red. Note that in *Trema* species NIN only occurs in truncated forms (Fig. 7). Included species: *Parasponia andersonii* (Pan); *Parasponia rigida* (Pri); *Parasponia rugosa* (Pru); *Trema orientalis* RG33 (Tor); *Trema orientalis* RG16 (TorRG16); *Trema levigata* (Tle); medicago (*Medicago truncatula*, Mt); lotus (*Lotus japonicus*, Lj); soybean (*Glycine max*, Glyma); peach (*Prunus persica*, Ppe); woodland strawberry (*Fragaria vesca*, Fvesca); black cotton poplar (*Populus trichocarpa*, Potri); eucalyptus (*Eucalyptus grandis*, Eugr); arabidopsis (*Arabidopsis thaliana*, At), jujube (*Ziziphus Jujube*) apple (*Malus × domestics*), mulberry (*Morns Notabilis*), hop (*Humulus Lupulus* (*natsume.shinsuwase.v1.0*)), and casuarina (*Casuarina glauca*). Node numbers indicate posterior probabilities, scale bar represents substitutions per site.

**Supplementary Figure 23:**
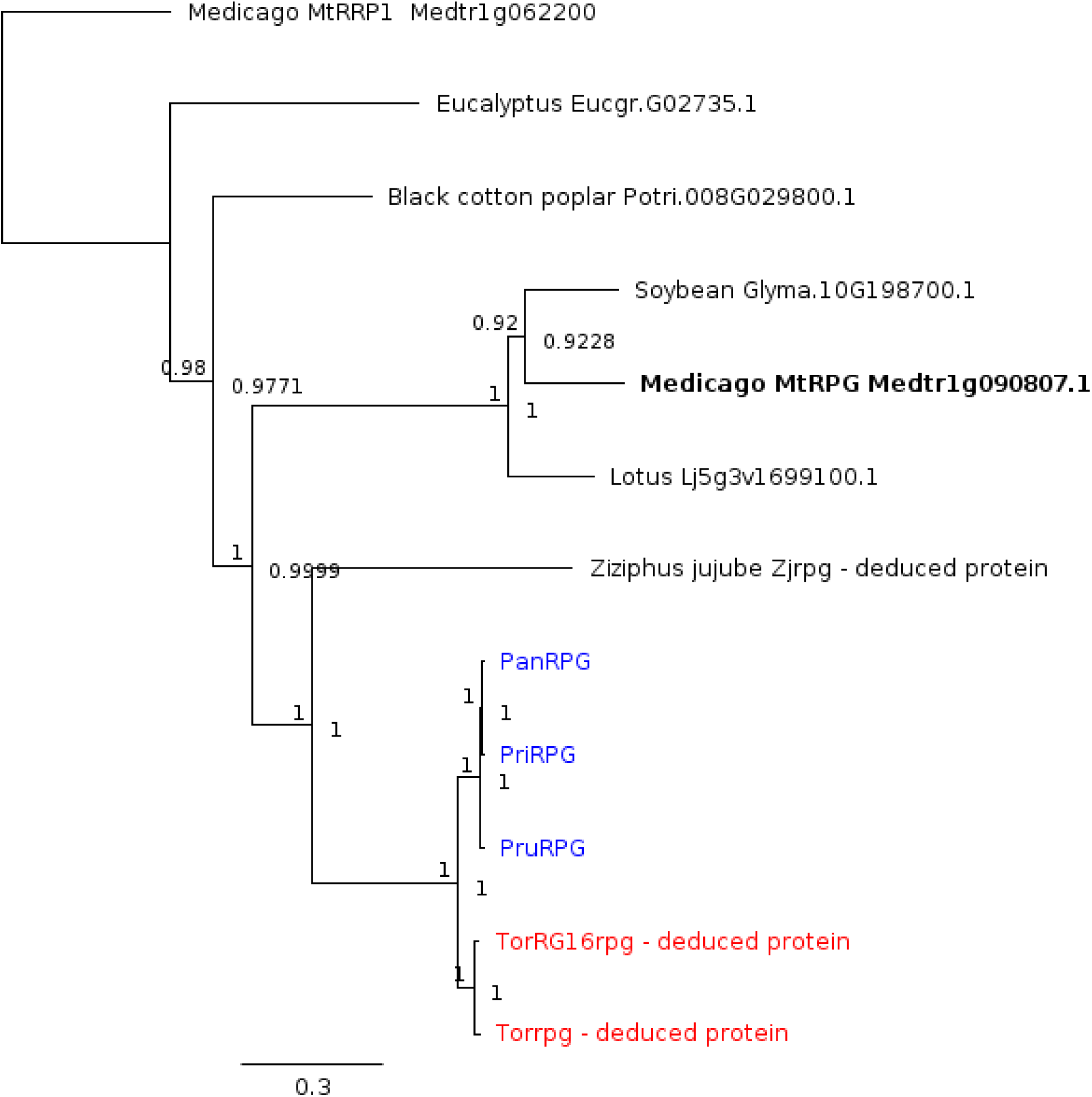
Phylogenetic reconstruction of the RPG orthogroup. Alignment of OG0014072 was supplemented with RPG homologs of additional species. *Parasponia* spp. marked in blue, Nitrogen fixation clade in bold. Included species: *Parasponia andersonii* (Pan) *Parasponia rigida* (Pri); *Parasponia rugosa* (Pru) *Medicago truncatula* (Mt); *Lotus japonicus* (Lj); *Glycine max* (Glyma), *Populus trichocarpa* (Potri); *Eucalyptus grandis* (Eugr). *Trema orientalis* RG33 (Tor); *Trema. orientalis* RG16 (TorRG16). *Ziziphus jujube* (Zj). No other functional RPG proteins could be detected in Rosales species, including *Fragaria vesca Ziziphus Jujube*, *Malus Domestica*, *Morus Notabilis*, and *Humulus Lupulus (natsume.shinsuwase.v1.0)*. Outgroup: *M. truncatula* MtRRP1 (RPG RELATED PROTEIN 1, Medtr1g062200.1). Node numbers indicate posterior probabilities, scale bar represents substitutions per site.

**Supplementary Figure 24:**
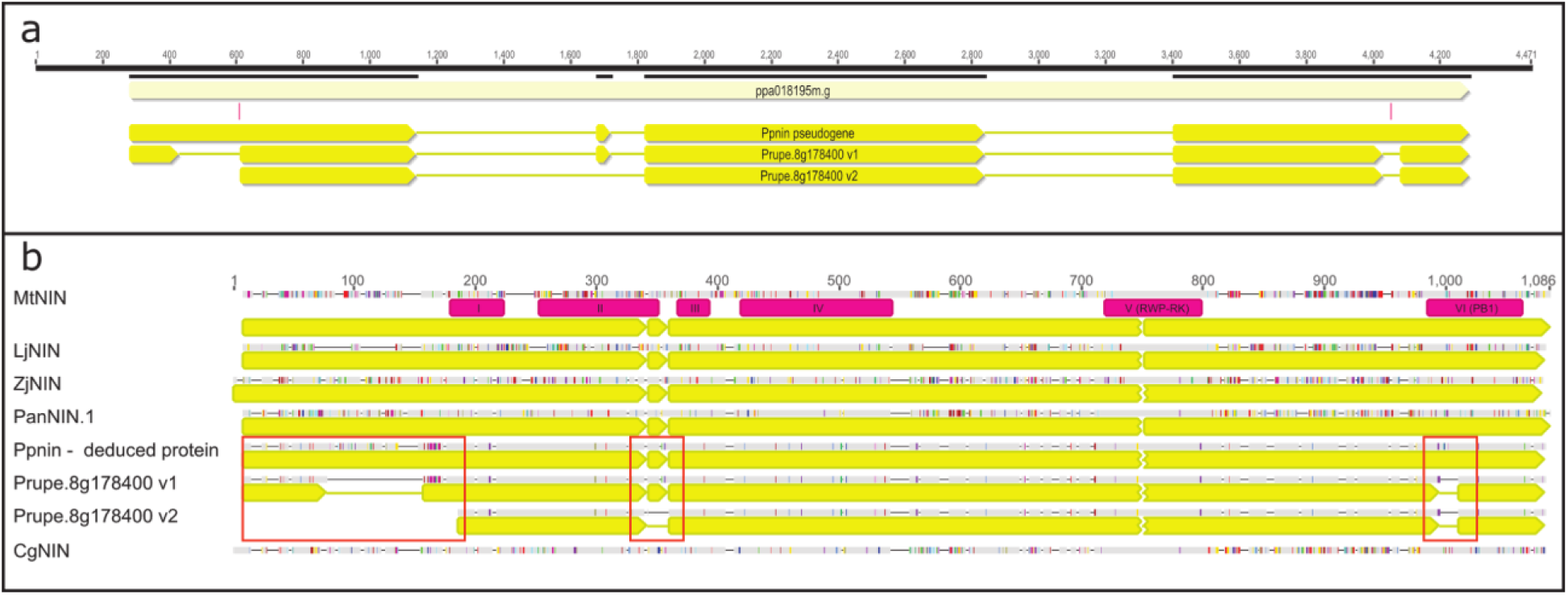
Annotation of *Prunus persica* locus ppa018195m.g representing *PpNIN*. **(a)** Comparison of the exon-intron structure of two publicly released gene models (named Prupe.8g17800_v1 and Prupe.8g178400_v2) and the gene model used here (*Ppnin* pseudogene). Yellow arrows: exons. Red bars indicate 2 single-nucleotide insertions that affect the coding region of the *Ppnin* pseudogene. **(b)** Alignment of derived/deduced NIN proteins of 3 *Prunus persica* gene models Prupe.8g17800_v1, Prupe.8g178400_v2, and *Ppnin* pseudogene, with *Medicago truncatula* MtNIN, *Lotus japonicus* LjNIN, *Ziziphus jujube* ZjNIN, *Parasponia andersonii* PanNIN.1, and *Casuarina glauca* CgNIN. Six conserved domains are annotated in MtNIN (cyan). Exon structure for all *NIN* genes indicated in yellow (except CgNIN for which no gene sequence is available). Deviations in the three *Prunus persica* derived/deduced NIN proteins are marked in red boxes.

**Supplementary Table 1:** *Parasponia andersonii* and *Trema orientals* RG33 putative orthologs of legume genes that function in rhizobium symbiosis.

Genes have been classified according to function of encoded proteins. CNV between *P. andersonii* and *T. orientalis* are marked in red. Genes for which no putative ortholog could be identified are indicated (not identified). The *P. andersonii* and *T. orientalis* genes are classified as either ‘putative ortholog’, ‘closest homolog’ or ‘inparalog’ depending on the phylogenetic relation with the legume symbiosis gene. Orthogroup number corresponds to orthogroups in Supplementary Table 7. It is indicated in case gene has been found to function in other symbiosis. AM: arbuscular mycorrhiza, and ANS: actinorhizal nodule symbiosis. Gm: *Glycine max;* Lj: *Lotus japonicus;* Ms: *Medicago sativa;* Mt: *Medicago truncatula;* Ps *Pisum sativum;* Pv: *Phaseolus vulgaris*.

**Supplementary Table 2:** Intergeneric crossings between *Parasponia* and *Trema* species.

Results column indicates whether intergeneric crosses could be obtained (positive) or not (negative).

**Supplementary Table 3:** *Parasponia*-*Trema* germplasm collection.

**Supplementary Table 4:** Genome size estimations based on estimated genome coverage.

**Supplementary Table 5:** Genome sequencing strategy.

**Supplementary Table 6:** Assembly results of *Parasponia* - *Trema* genome sequences. #N is number of gap sequences; GC% is guanine-cytosine content; BUSCO^87^ and CEGMA^86,87^ are tools that assess completeness of genome assemblies by checking sets of conserved genes. For BUSCO a set of 1,440 plant specific genes was used.

**Supplementary Table 7:** Inferred orthogroups.

Eurosid orthogroups generated by OrthoFinder based on gene models from *Parasponia andersonii*, *Trema orientalis*, *Medicago truncatula*, *Glycine max*, *Fragaria vesca*, *Populus trichocarpa*, *Arabidopsis thaliana* and *Eucalyptus grandis*.

**Supplementary Table 8:** Gene models in *Parasponia andersonii* and *Trema orientalis* RG33 reference genomes.

Inparalogs: species specific duplications; singletons: loss of gene in other species; multi-orthologs: duplication in the other species; CNVs: copy number variants. We found no significant enrichment of these CNVs in the symbiosis genes in Supplementary Table 1 and nodule enhanced genes in Suppplementary table 9 (hypergeometric test, p = 0.99). For BUSCO a set of 1,440 plant specific genes was used.

**Supplementary Table 9:** *Parasponia* nodule-enhanced gene set.

*P. andersonii* genes with enhanced expression in three nodule developmental stages compared to non-inoculated roots (>2-fold increase). Stage 1: initial stages of colonization when infection threads entering the host cells. Stage 2: progression of rhizobium infection in nodule host cells, Stage 3: nodule cells filled with fixation threads. Plants were inoculated with *M. plurifarium* BOR2. OrthoGroup number corresponds to Supplementary Table 7, STAT: trident alignment conservation score, CLASS: gene homology classification (nv = orthologous pair not validated by whole-genome alignments) FC: gene expression log fold-change in nodule versus root, P: P-value adjusted for multiple testing based on false discovery rate estimation. Genes that are putatively orthologous to legume genes with symbiotic function are classified as ‘LEGUME SYMBIOSIS GENE’. Conserved CNVs between *Parasponia* and *Trema* species are shaded pink.

**Supplementary Table 10:** Commonly utilized symbiosis gene set.

*P. andersonii* genes with enhanced expression in three nodule developmental stages compared to non-inoculated roots (>2-fold increase) (Supplementary Table 9), of which a putative ortholog in *Medicago truncatula* also has been identified as a gene with a nodule enhanced expression^31^. Cluster numbers indicate expression profile clusters of commonly utilized genes as shown in Fig. 5. DESeq2 normalized read counts are included for *P. andersonii* and hybrid roots, hybrid nodules and three stages of *P. andersonii* nodules. Stage 1: initial stages of colonization when infection threads enter the host cells. Stage 2: progression of rhizobium infection in nodule host cells, Stage 3: nodule cells filled with fixation threads. Plants were inoculated with *M. plurifarium* BOR2. OrthoGroup number corresponds to Supplementary Table 7, STAT: trident alignment conservation score, CLASS: *Parasponia*-*Trema* gene homology classification (nv = orthologous pair not validated by whole-genome alignments), FC: gene expression log fold-change gene expression in nodule versus roots, P: P-value adjusted for multiple testing based on false discovery rate estimation. Genes that are putatively orthologous to legume and actinorhizal genes with symbiotic function are classified as ‘LEGUME SYMBIOSIS GENE’. Conserved CNVs between *Parasponia* and *Trema* spp are in bold.

**Supplementary Table 11:** Sequenced RNA samples.

**Supplementary Table 12:** GenBank Accession numbers of sequences.

Tab 1: sequences used in phylogenetic reconstructions of Cannabaceae. Tab 2: sequences of genes with copy number variants. Sequences generated for this study are in bold; pseudogenes are marked with grey background.

**Supplementary Data File 1: Phylogenetic analysis of *Parasponia andersonii* and *Trema orientalis* RG33 putative orthologs of legume genes that function in symbioses.** Phylogenetic trees (based on Neighbour Joining) of orthogroups containing published genes that function in symbiosis (see also Supplementary Table 1). OrthoGroup number corresponds to Supplementary Table 7; Node labels indicate bootstrap support values. Legume symbiosis genes and symbiotic homologs from actinorhizal species are marked in bold; proteins from *Arabidopsis thaliana* (AT) are marked in green; *Eucalyptus grandis* (Eucgr) in olive; *Populus trichocarpa* (Potri) in light blue; *Medicago truncatula* (Medtr) in purple; *Glycine max* (Glyma) in mint; *Fragaria vesca* (Fvesca) in pink; *P. andersonii* (Pan) in dark blue; and *T. orientalis* (Tor) in dark red. Legume or actinorrhizal symbiosis genes from species not included in the orthogroup inferences are in black. Agl: *Alnus glutinosa;* Cgl: *Casuarina glauca;* Dgl *Datisca glomerata;* Lja: *Lotus japonicus;* Msa: *Medicago sativa;* Mtr: *Medicago truncatula;* Phy: *Petunia hybrida;* Psa: *Pisum sativum;* Pvu: *Phaseolus vulgaris*.

**Supplementary Data File 2: Phylogenetic analysis of genes utilized in *Parasponia* and medicago root nodules.** Phylogenetic trees (based on Neighbour Joining) of orthogroups containing genes with significantly enhanced expression level in any of three *Parasponia* nodule developmental stages (Supplementary Fig. 11; Supplementary Table 9) as well as genes with significantly enhanced expression in nodules of medicago^31^ (Supplementary Table 10). OrthoGroup number corresponds to Supplementary Table 7; Node labels indicate bootstrap support values. Nodule-enhanced genes are marked in bold; proteins from *Arabidopsis thaliana* (AT) are marked in green; *Eucalyptus grandis* (Eucgr) in olive; *Populus trichocarpa* (Potri) in light blue; *Medicago truncatula* (Medtr) in purple; *Glycine max* (Glyma) in mint; *Fragaria vesca* (Fvesca) in pink; *P. andersonii* (Pan) in dark blue; and *T. orientalis* (Tor) in dark red. Agl: *Alnus glutinosa;* Cgl: *Casuarina glauca;* Dgl *Datisca glomerata;* Lja: *Lotus japonicus;* Msa: *Medicago sativa;* Mtr: *Medicago truncatula;* Phy: *Petunia hybrida;* Psa: *Pisum sativum;* Pvu: *Phaseolus vulgaris*.

